# Systemic delivery of a CXCR4-CXCL12 signaling inhibitor encapsulated in synthetic protein nanoparticles for glioma immunotherapy

**DOI:** 10.1101/2021.08.27.457953

**Authors:** Mahmoud S Alghamri, Kaushik Banerjee, Anzar A Mujeeb, Ayman Taher, Rohit Thalla, Brandon L McClellan, Maria L Varela, Svetlana M Stamatovic, Gabriela Martinez-Revollar, Anuska Andjelkovic-Zochowska, Jason V Gregory, Padma Kadiyala, Alexandra Calinescu, Jennifer A Jiménez, April A Apfelbaum, Elizabeth R Lawlor, Stephen Carney, Andrea Comba, Syed Mohd Faisal, Marcus Barissi, Marta B. Edwards, Henry Appelman, Michael R. Olin, Joerg Lahann, Pedro R. Lowenstein, Maria G. Castro

**Author notes:** Joint First Authors (Mahmoud S Alghamri, Kaushik Banerjee, Anzar A Mujeeb).

## Abstract

Glioblastoma multiforme (GBM) is an aggressive primary brain tumor, with poor prognosis. Major obstacles hampering effective therapeutic response in GBM are tumor heterogeneity, high infiltration of immunosuppressive myeloid cells, and the presence of the blood-brain barrier. The C-X-C Motif Chemokine Ligand 12/ C-X-C Motif Chemokine Receptor 4 (CXCL12/ CXCR4) signaling pathway is implicated in GBM invasion and cell cycle progression. While the CXCR4 antagonists (AMD3100) has a potential anti-GBM effects, its poor pharmacokinetic and systemic toxicity had precluded its clinical application. Moreover, the role of CXCL12/ CXCR4 signaling pathway in anti-GBM immunity, particularly in GBM-mediated immunosuppression has not been elucidated. Here, we developed a synthetic protein nanoparticle (SPNPs) coated with the cell-penetrating peptide iRGD (AMD3100 SPNPs) to target the CXCR4/CXCL12 signaling axis in GBM. We showed that AMD3100 SPNPs effectively blocked CXCR4 signaling in mouse and human GBM cells *in vitro* as well as in GBM model *in vivo*. This results in inhibition of GBM proliferation and induction of immunogenic tumor cell death (ICD) leading to inhibition of GBM progression. Our data also demonstrate that blocking CXCR4 sensitizes GBM cells to radiation, eliciting enhanced release of ICD ligands. Combining AMD3100 SPNPs with radiotherapy inhibited GBM progression and led to long-term survival; with 60% of mice remaining tumor-free. This was accompanied by an anti-GBM immune response and sustained immunological memory that prevented tumor recurrence without further treatment. Finally, we showed that systemic delivery of AMD3100 SPNPs decreased the infiltration of CXCR4^+^ monocytic myeloid-derived suppressor cells to the tumor microenvironment. With the potent ICD induction and reprogrammed immune microenvironment, this strategy has significant potential for future clinical translation.

**Graphical abstract:** Immunological mechanism targeting Glioblastoma (GBM) upon blocking CXCR4 signaling pathway with AMD3100-conjugated nanoparticles (SPNPs).
(1) Radiotherapy induces glioma cell death, followed by Damage-associated molecular patterns (DAMPs) release. Dendritic cells (DC) are activated by DAMPs and migrate to the regional lymph node where they prime cytotoxic T lymphocyte immune response. Tumor-specific cytotoxic T cells infiltrate the tumor and attack glioma cells. (2) Glioma cells express CXCR4, as well its ligand CXCL12. CXCL12 induces glioma cell proliferation and, (3) as well as mobilization in the bone marrow of CXCR4 expressing myeloid MDSC, which will infiltrate the tumor, and inhibit tumor-specific cytotoxic T cells activity. GEMM of glioma when treated systemically with SPNPs AMD3100 SPNPs plus radiation, nanoparticles block the interaction between CXCR4 and CXCL12, thus (4) inhibiting glioma cell proliferation and (5) reducing mobilization in the bone marrow of CXCR4 expressing myeloid MDSC, (6) generating a reduced MDSC tumor infiltration, as well as releasing MDSC inhibition over tumor specific cytotoxic T cell response.

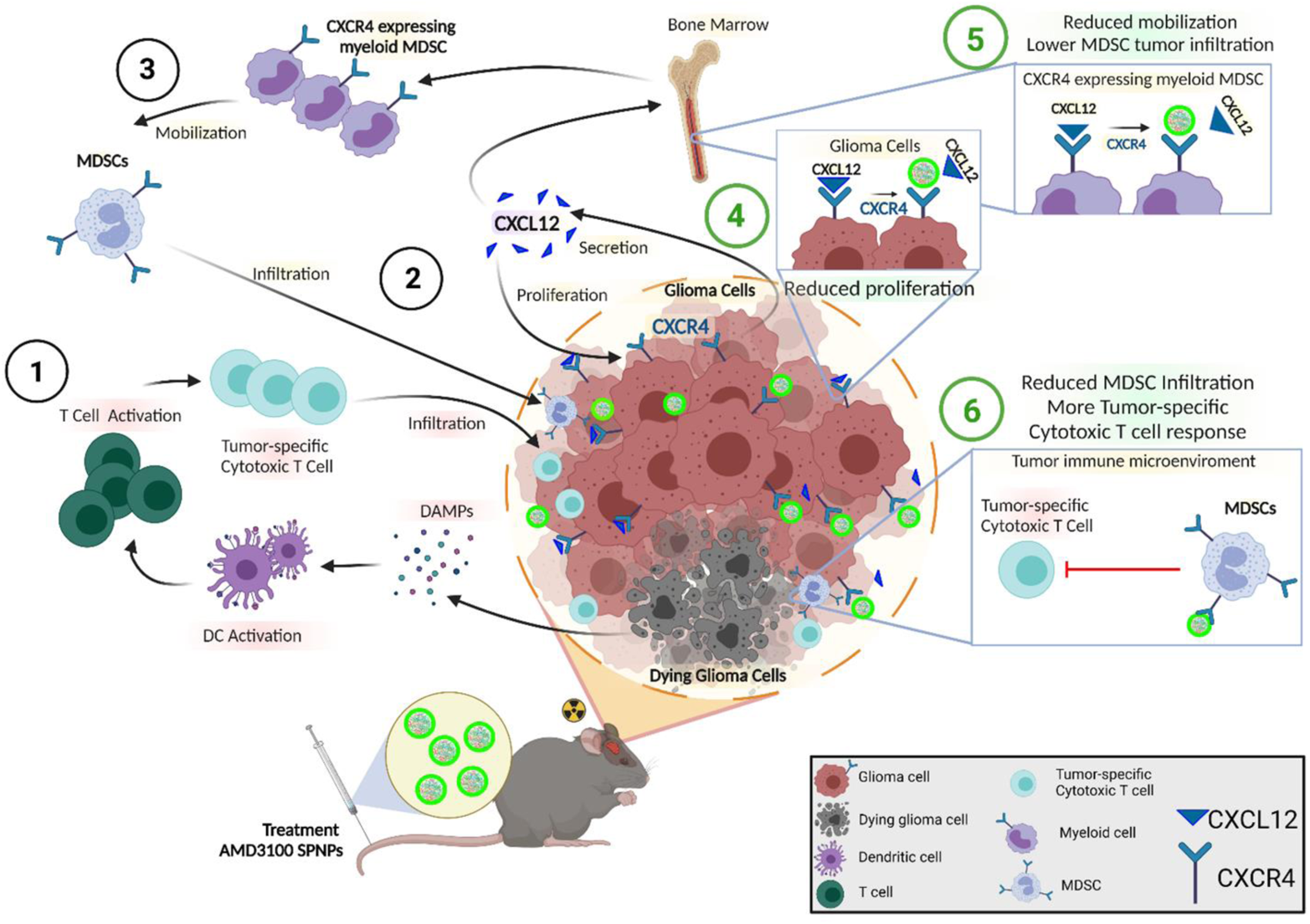

## Introduction

Glioblastoma (GBM) is an aggressive brain tumor with poor prognosis, characterized by a high level of cellular and molecular heterogeneity, high proliferative capacity, and invasive borders, making GBM challenging to treat ^1, 2^. The invasive characteristics of GBM lead to infiltration of tumor cells into the normal brain tissue, making GBM difficult to completely resect and increasing the likelihood of tumor recurrence. This is compounded by the presence of immunosuppressive immune cells which hinder the effectiveness of immunotherapies ^3, 4^.

Moreover, the presence of the blood-brain barrier (BBB) provides both physical and biochemical barriers to drug delivery into the brain ^5, 6^. This limits the brain permeability of many chemotherapeutic drugs, including monoclonal antibodies, antibody-drug conjugates, and hydrophilic molecules that do not readily cross lipid bilayers ^7^.

The CXCL12/CXCR4 signaling pathway is involved in multiple physiological processes including hematopoiesis ^8^, retention of hematopoietic stem cells (HSCs) in the bone marrow, and central nervous system (CNS) development ^9, 10^. Several studies have illustrated the involvement of activated CXCL12/ CXCR4 signaling in solid cancers in promoting survival, growth, and metastasis ^11–14^. In GBM, it has been previously demonstrated the CXCL12/CXCR4 signaling pathway is important for sustained invasion ^15^, enhanced angiogenesis ^16^, and maintenance of glioma stem-cell migration and therapeutic resistance ^17^. This pathway is particularly upregulated under hypoxic conditions, a feature that is associated with worse prognosis in GBM ^18–21^.

CXCR4 is expressed by many cells, including hematological progenitor cells, myeloid cells, stromal fibroblasts, endothelial cells, epithelial cells, and tumor cells ^22–24^. Most importantly, CXCR4 expression on myeloid cells promotes trafficking of myeloid-derived suppressor cells (MDSCs) in several cancers such as osteosarcoma ^25^, ovarian cancer ^26^, colorectal cancer ^27^, metastatic melanoma ^28^, and leukemia ^29^. Enhanced MDSCs infiltration promotes an immunosuppressive tumor microenvironment and further contributes to immunotherapeutic resistance. Thus, blocking CXCR4 provides an attractive target to promote effective immunotherapy. However, the impact of CXCL12/CXCR4 signaling on immune-mediated therapeutic outcomes in GBM and its impact on reprograming the immunosuppressive tumor immune microenvironment (TIME) has not yet been elucidated.

The presence of the BBB limits the efficacy of both conventional and novel therapies for GBM. Thus, the development of nano-based therapies that have the ability to cross the BBB is of great interest to improve GBM clinical outcomes. A wide variety of nanoparticles have been tested in preclinical settings to facilitate effective drug delivery into brain tumors ^30–33^. Remaining challenges in nanoparticle safety, include the fact that most of the widely used NPs are composed of non-organic materials that tend to accumulate in the liver ^34–36^. Another compounding factor, especially for GBM therapeutics, relates to their limited BBB penetration capacity ^34–36^. We recently developed biologically compatible NPs comprised of polymerized human serum albumin (HSA) and oligo ethylene glycol (OEG), equipped with the cell-penetrating peptide iRGD ^37^. HSA was used as the primary component because of its biological compatibility, and well-studied kinetic, and interactions with other cellular proteins and receptors ^38, 39^. The incorporation of the tumor-targeting-penetrating peptide, iRGD, results in the ability of the SPNPs to target the tumor mass, after systemic delivery ^37^. We previously demonstrated that these SPNPs loaded with siRNA against signal transducer and activator of transcription 3 (STAT3i) are able to silence STAT3 within the GBM cells *in vivo* ^37^. The main advantage of the SPNPs is that they are able to deliver their therapeutic cargo to the target tissue (GBM) after systemic delivery.

Here, we used SPNPs loaded with the CXCR4 inhibitor, AMD3100, to block CXCR4 signaling in an aggressive intracranial GBM model. We demonstrate that blocking CXCR4 results in reduced MDSCs trans-endothelial migration *in vitro* and reduced infiltration of CXCR4^+^ M-MDSCs to the GBM TIME *in vivo*. Interestingly, we showed that blocking CXCR4 sensitizes GBM cells to radiation-induced immunogenic cell death (ICD) which triggers an anti-GBM adaptive immune response. With the potent ICD induction and reprogrammed immunosuppressive microenvironment, SPNPs-elicited antigen presentation, immune priming, and GBM specific immunological T-cell mediated immunity.

## Results

### Infiltration of CXCR4^+^ M-MDSCs within the tumor immune microenvironment in genetically engineered mouse GBM models

To study the impact of CXCL12/CXCR4 signaling pathway on the glioma infiltrating myeloid cells to the glioma tumor microenvironment, we used three different implantable glioma models. Two models were aggressive, fast-growing tumors developed using the SB-generated neurospheres, i.e. OL61, and RPA as described before (Figure 1A) ^40–42^. The third model was a less aggressive, slower-growing GBM model developed using RCAS-TVA system (Arf^−/−^) (Figure 1A) ^40^. When comparing the median survival (MS) between the three groups, we found that both OL61 and RPA had a median survival (MS= 18dpi and 14dpi, respectively). Whereas the Arf^−/−^ tumor-bearing mice had a longer MS compared to the other groups (Arf^−/−^ MS= 69dpi, P< 0.001) (Figure 1B). Next, we investigated the frequencies and the subsets of CXCR4^+^ MDSCs infiltrating in each GBM model. MDSCs infiltrating the GBM could be of granulocytic (PMN-MDSCs) (CD45^high^/CD11b^+^/Ly6G^+^/Ly6C^low^) or monocytic (M-MDSCs) (CD45^high^/CD11b^+^/Ly6G^-^/Ly6C^high^) origin. First, we characterized the MDSCs subsets infiltrating each GBM model and found that in contrast to PMN-MDSCs which showed no differences in frequencies between the three groups, the frequencies of M-MDSCs infiltrating the tumors in the aggressive GBM models (i.e. OL61 and RPA) were significantly higher compared to the Arf^−/−^ model (36% and 35% vs 19%, respectively, (P<0.01, Figure 1C-E)). Moreover, we found that the majority of CXCR4 expressing MDSCs belong to M-MDSCs, with no difference in CXCR4^+^ PMN-MDSCs between the three tumor models (Figure 1F-I). This suggests that the frequency of CXCR4^+^ M-MDSCs which infiltrate GBM correlates with tumor aggressiveness.

**Figure 1.**
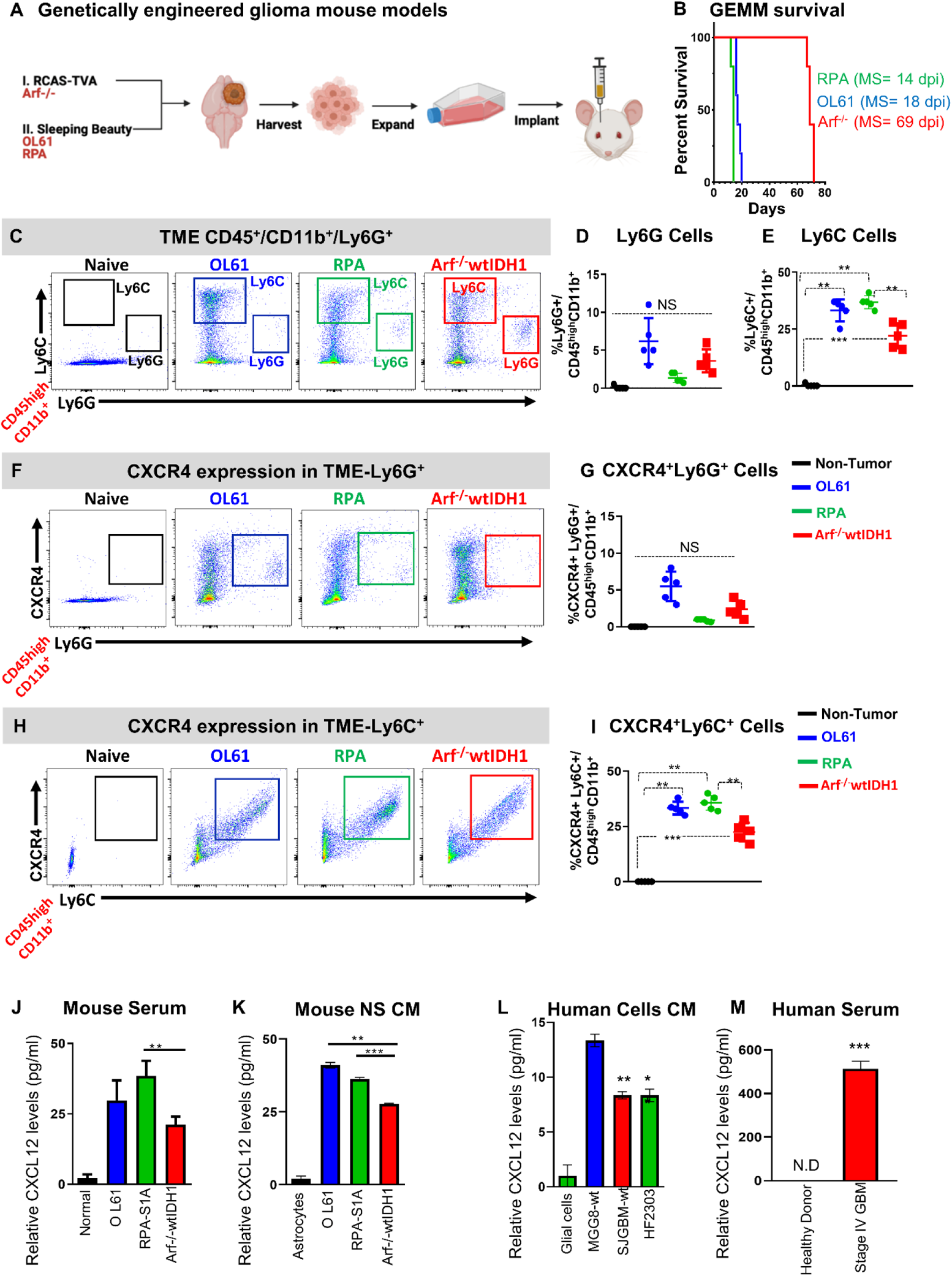
Aggressive genetically engineered GBM models are associated with activated CXCR4/CXCL12 signaling and infiltrating of immunosuppressive myeloid cells. **(A)** Experimental design of the generation of GBM models using the sleeping beauty (SB, aggressive) and RCAS-TVA technology (slow-growing). Neurospheres from each model were harvested, cultured and used for intracranial implantation in animals. (B) Kaplan-Meier survival curves of mice bearing RCAS-TVA, OL61, or RPA (MS: median survival). (C-E) Characterization of granulocytic and monocytic myeloid cell population (Ly6G vs Ly6C) in normal brain or OL61, RPA, and Arf^−/−^ tumor bearing mice. Arf^−/−^ wtIDH1 tumor bearing mice display lower percentage of monocytic myeloid cells (M-MDSCS; Ly6C+) compared to OL61 and RPA tumor bearing mice. (F, G) Flow analysis of the Ly6G+ CXCR4+ myeloid cells in normal brain or in the tumor from OL61, RPA, and Arf^−/−^ implanted mice. (H-I) Flow analysis of the Ly6C+ CXCR4+ myeloid cells in normal brain or in the tumor from OL61, RPA, and Arf^−/−^ implanted mice. Quantitative ELISA of the CXCL12 level in mouse serum of tumor-bearing animals (J), conditioned media from cultured mouse, (K) conditioned media from human cells (L), and serum from control and GBM patients (M). *P<0.05, \**\* *P<0.01, *** P<0.001.* ANOVA, (N=5/group).

CXCL12 is the primary ligand for CXCR4 receptors expressed by GBM cells, and its expression is responsible for the homeostasis of HSCs in the BM ^10, 17, 43^. We hypothesized that the enhanced infiltration of M-MDSCs in aggressive GBM models results from activation of CXCL12/CXCR4 signaling. Quantitative ELISA analysis showed that the level of CXCL12 was lower in the Arf^−/−^ GBM model in both mouse serums from implanted animals as well as conditioned media of cultured GBM cells (P<0.01, Figure 1J, K). We also observed that CXCL12 is expressed at higher levels in patient-derived cells (MGG8, SJGBM, and HF2303) as well as in serum from stage IV GBM patients compared to serum from healthy donors (10-fold increase vs control) (Figure 1L, M). Overall, these results suggest that CXCL12/CXCR4 signaling is activated in aggressive GBM and is associated with enhanced infiltration of immunosuppressive M-MDSCs.

### CXCR4 is expressed primarily by monocytic MDSCs in the spleen and blood from GBM mouse models and is associated with poor prognosis in human GBM patients

We next asked if the change we observed in the frequency of CXCR4^+^ MDSCs in the TIME from different tumor models is associated with changes in CXCR4 expressing myeloid cells in blood, spleen, and BM. Our results show that in both the blood and spleen, PMN-MDSCs were the dominant population with no difference in the frequencies between the three tumor models (Figure 2A, B, D, E). However, the frequency of M-MDSCs was significantly higher in the blood from OL61 and RPA tumor-bearing mice compared to the frequency observed in Arf^−/−^ tumor-bearing mice (Figure 2A, B). Like the TIME, we found that there was an increased expression of CXCR4 by the M-MDSC population within OL61 tumors compared to the two other models (Figure 2C). There were no differences in the frequencies of M-MDSC and PMN-MDSC populations in the spleen of the three GBM models, however, M-MDSCs were the major population expressing CXCR4 (Figure 2D-F).

**Figure 2.**
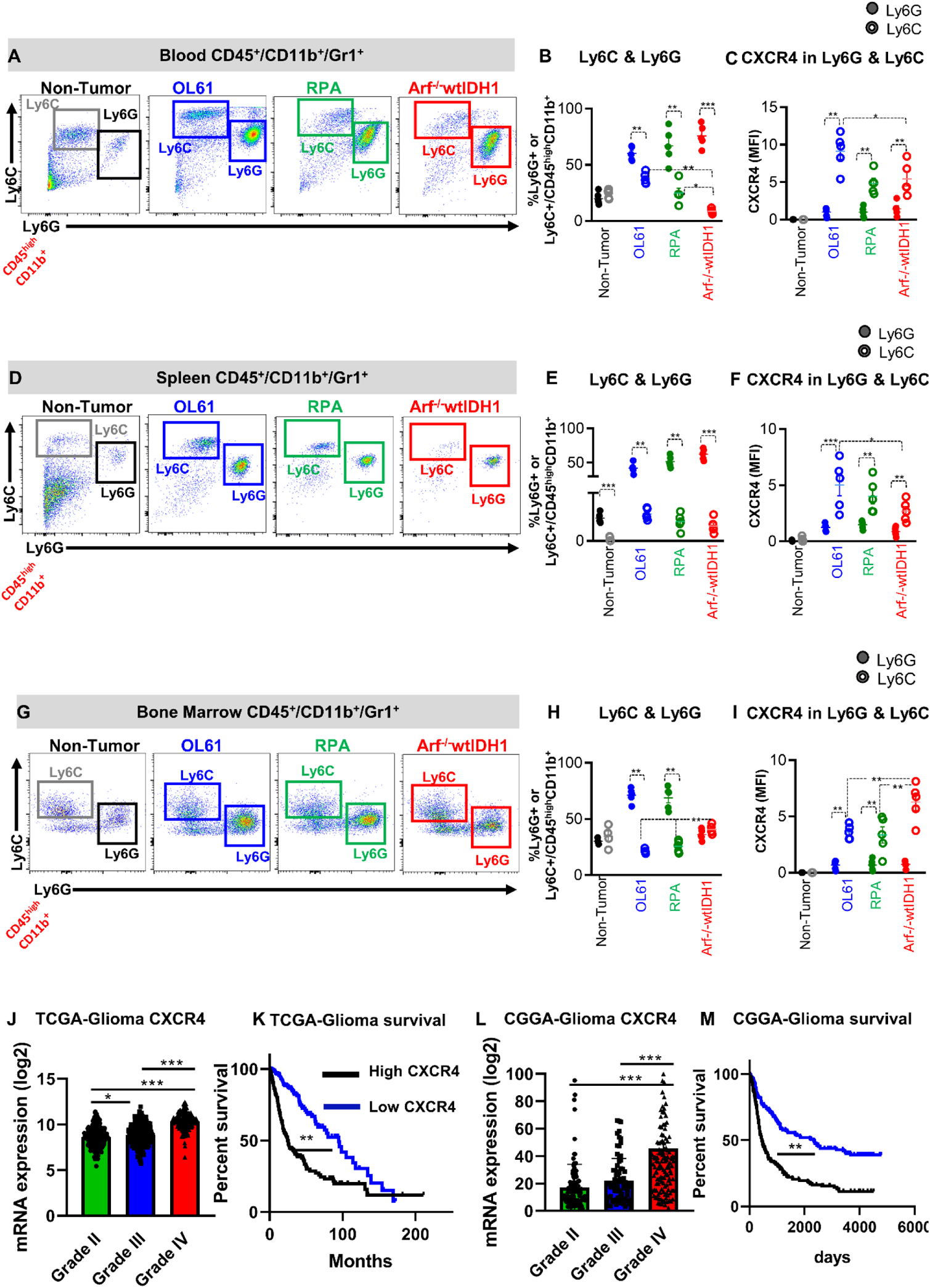
CXCR4 is expressed primarily by monocytic MDSCs (CD45^high^/CD11b^+^/Ly6C^high^) and is associated with poor prognosis. **(A, B)** Representative flow cytometry plots and quantification of the percentage of PMN-MDSCs (CD45^high^/CD11b^+^/Ly6G^+^/Ly6C^low^) or M-MDSCs (CD45^high^/CD11b^+^/ Ly6C^high^) in bone marrow (BM) from normal mice (N), and mice implanted with OL61, RPA, or Arf^−/−^ wtIDH1 neurospheres. **(C)** Quantitative analysis of CXCR4 expression in conditions from **(B)**. **(D, E)** Representative flow cytometry plots and quantification of the percentage of PMN-MDSCs (CD45^high^/CD11b^+^/Ly6G^+^/Ly6C^low^) or M-MDSCs (CD45^high^/CD11b^+^/ Ly6C^high^) in blood from normal mice (N), and mice implanted with OL61, RPA, or Arf^−/−^ wtIDH1 neurospheres. **(F)** Quantitative analysis of CXCR4 expression in conditions from **(E)**. **(G, H)** Representative flow cytometry plots and quantification of the percentage of PMN-MDSCs (CD45^high^/CD11b^+^/Ly6G^+^/Ly6C^low^) or M-MDSCs (CD45^high^/CD11b^+^/ Ly6C^high^) in spleen from normal mice (N), and mice implanted with OL61, RPA, or Arf^−/−^ wtIDH1 neurospheres. **(I)** Quantitative analysis of CXCR4 expression in conditions from **(H)**. **(J)** Analysis of *CXCR4* gene expression for glioma patients according to their grade, Grade II (n=226), Grade III (n= 244), and Grade IV (n=150). **(K)** Kaplan-Meier survival analysis of TCGA glioma patients with high vs low level of *CXCR4* expression. **(L)** Analysis of *CXCR4* gene expression for glioma patients in CGGA database according to their grade, Grade II (n=103), Grade III (n= 79), and Grade IV (n=139). **(M)** Kaplan-Meier survival analysis of CGGA glioma patients with high vs low level of *CXCR4* expression. * *P<0.05, \**\* *P<0.01, \**\*\* *P<0.005,* One-way ANOVA, (N=5/group).

On the other hand, we found that the frequency of M-MDSCs was higher in the BM from the Arf^−/−^ GBM mouse model when compared to Ol61 and RPA GBM models (Figure 2G, H). This corresponded to higher expression of CXCR4 in the M-MDSCs residing in the BM of the Arf^−/−^ GBM model (Figure 2I). These results suggest that there is more trafficking of CXCR4^+^ M-MDSCs from the bone morrow into circulation, and to the TIME, in the OL61 and RPA GBM models, that promote an immunosuppressive GBM TIME.

To evaluate if CXCR4 expression is associated with GBM tumor progression in clinical settings, we queried the TCGA data and analyzed *CXCR4* expression in three different glioma grades (Grade II, III, and IV). We also examined the MS of glioma patients in the context of CXCR4 (*CXCR4*) expression (i.e. MS of patients expressing high vs low levels of *CXCR4*). TCGA-Glioma analysis revealed that *CXCR4* gene expression level is correlated with tumor grade, and its expression is associated with a worse prognosis in glioma patients (Figure 2J, K). Similar results were found when analyzing data from the Chinese Glioma Genome Atlas (CGGA) (Figure 2L, M). These results suggest that there is a unique correlation between CXCR4 expression and prognosis in glioma patients.

### CXCR4 signaling disrupts brain endothelial cells’ barrier permeability and enhances myeloid cells’ infiltration

To assess the effect of CXCR4 signaling on blood-brain barrier (BBB) integrity and permeability, we used an *in vitro* model for the BBB established in a transwell dual-chamber system. In this assay, brain microvascular endothelial cells (BECs) were grown to confluence in the upper chamber of collagen type IV coated filters in a trans-well dual-chamber system. To mimic the BBB biology, pericytes were grown on the opposite side (basolateral) of the filter (Figure 3A). First, we tested the effect of CXCR4 inhibition on myeloid cells’ transmigration from the apical-basolateral side. Conditioned media (CM) collected from OL61, Arf^−/−^ and RPA cultures treated with either vehicle or the CXCR4 inhibitor AMD3100 (according to the IC_50_ (Figure S1A)), were placed at the basolateral side (Figure 3A). MDSCs were placed on the apical side of the BBB transwell system. Results showed that treatment with AMD3100 significantly reduced the number of myeloid cells transmigrated through the basolateral chamber in all three GBM cell lines tested (Figure 3B).

**Figure 3.**
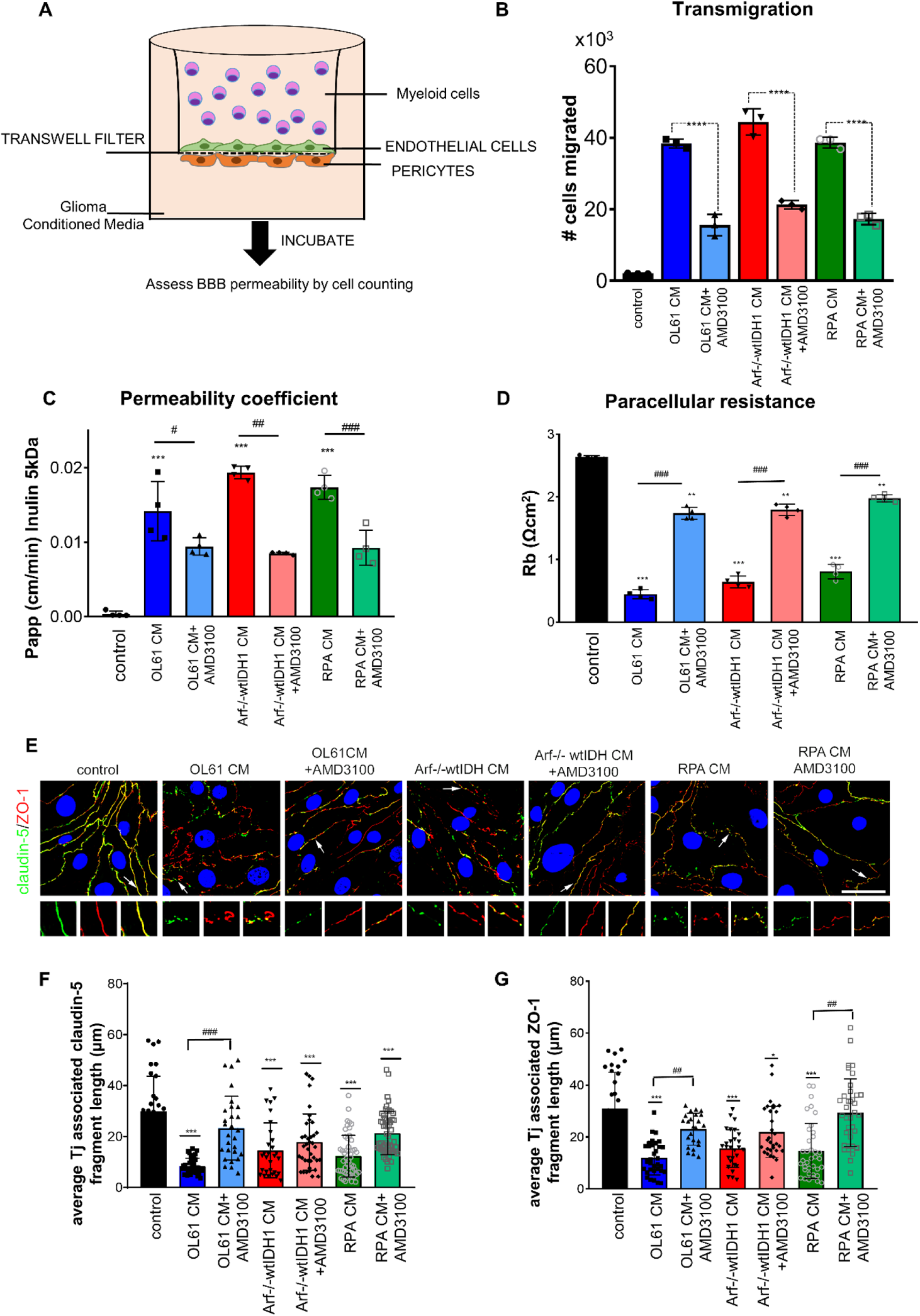
CXCR4 signaling enhances myeloid cells transmigration and increases brain endothelial cell (BEC) barrier permeability. **(A**) Diagram of transwell dual-chamber system used for cell migration assay. **(B)** Bar graph represents the number of the myeloid cells migrated through the endothelial-pericytes transmembrane. **(C)** Permeability coefficient (Papp) for FITC-inulin in mBMEC monolayers exposed to condition media collected from OL61, Arf^−/−^wtIDH and RPA cells with or without CXCR4 inhibitor AMD3100 for 24 hrs. **(D)** Bar graph represents paracellular resistance (Rb) value at 24 hrs for all analyzed groups. **(E)** Immunofluorescence staining for tight junction (Tj) proteins claudin-5 and ZO-1 in control and cells exposed to OL61, OL61+AMD3100, Arf^−/−^wtIDH, Arf^−/−^wtIDH+AMD3100, and RPA and RPA+AMD3100 for 24 hrs. Arrow and magnified images indicate pattern and colocalization of claudin-5 and ZO-1 on the cell border. Scale bar 50mm. Quantitation of the average TJ-associated **(F)** claudin-5 and **(G)** ZO-1 fragment length in claudin-5/ZO-1 costained immunofluorescent images in control and cells exposed to OL61, OL61+AMD3100, Arf^−/−^wtIDH, Arf^−/−^wtIDH+AMD3100, and RPA and RPA+AMD3100 for 24 hrs. Data are shown as means ± SD. *n* = 3-5; ***p>0.0001 and **p>0.001 comparing to control. ###p>0.0001 comparing experimental groups with and without inhibitor AMD3100.

We also analyzed the effects of conditioned media (CM) collected from glioma cells on the barrier properties of the mouse brain endothelial cells (mBECs). Results showed that CM collected from OL61, RPA and Arf^−/−^ GBM cells increased brain endothelial barrier permeability for the small molecular weight tracers, i.e., Cadaverine (1kDa), Dextran (3kDa), and Inulin (5kDa), but not for high molecular weight Dextran (10kDa) (Figure S2, and Figure 3C). Our results also show that blocking CXCR4 with AMD3100 rescued the impaired barrier integrity (Figure S2, and Figure 3C). This was validated in multi-frequency measurements of barrier properties (resistance, capacitance, and impedance) showing that AMD3100 treatment caused a two-fold increase in the paracellular resistance (Figure 3D). Morphologically, the effects of GBM CM on brain endothelial barrier integrity were associated with alteration in tight junction (Tj) protein expression, localization, and complex organization (Figure 3E). Because of their role in preserving the integrity of the BBB ^44^, the major occlusion Tj protein claudin-5 and ZO-1 were analyzed by immunofluorescence and confocal microscopy. Results showed that conditioned media collected from the three cultured GBM cells caused defragmented, and punctate staining of claudin-5 and ZO-1 in mBECs (Figure 3E-G). This was associated with dislocation of both Tj proteins from mBECs and significant reduction of claudin-5 and ZO-1 associated Tj length (Figure 3E-G). We confirmed the decreased claudin-5 and ZO-1 protein expression levels by western blotting (Figure S3). As protein expression only partially characterizes the alteration in the Tj complex, we took one step further and analyzed the interactions between the ZO-1 and claudin-5 by performing a proximal ligation assay (PLA) to detect protein-protein interactions at higher specificity and sensitivity. Consistent with the immunofluorescence, PLA analysis showed decreased interactions between claudin-5 and ZO-1 in mBECs treated with CM only (P<0.01, Figure S4). Blocking CXCR4 resulted in more than 2-fold increase in the interactions between claudin-5 and ZO-1 (P<0.01, Figure S4). This functional rescue of the barrier upon CXCR4 blockade was accompanied by re-establishing the Tj complex, shown by restoration of claudin-5 and ZO-1 localization on the cell border mired in increased length of the Tj fragment and increased the number of interactions between claudin and ZO-1 (P<0.01, Figure 3F, G, Figure 3S, Figure S5). We further validated this effect using human brain endothelial cells (hBECs) incubated with CM collected from two patient-derived GBM cells (SJGBM2 and HF2303) in the presence or absence of either vehicle or AMD3100 (according to the IC_50_ (Figure S1B)). Consistent with the mouse data, our results showed that blocking CXCR4 expression reversed the effect of CM-mediated disruption of ZO-1 and claudin-5 (P<0.01, Figure S5). These results verified the protective effects of the CXCR4 blockade from GBM-mediated derangement of the BBB integrity.

### Blocking of CXCR4 sensitizes mouse and human glioma cells to radiotherapy and induces immunogenic cell death

CXCR4 signaling is crucial for GBM cell survival and proliferation ^17, 43^. We sought to determine whether CXCR4 inhibition would induce radio-sensitization and also induce immunogenic cell death (ICD). First, we performed cell viability and real-time cell proliferation assay in three mouse glioma cell lines (i.e. RPA, OL61, Arf^−/−^) and three human glioma cell lines (i.e. MGG8, SJGBM2, HF2303) treated with either saline, AMD3100, IR, or AMD3100+ IR (Figure 4A). Results showed that treatment with either AMD3100 or IR alone markedly inhibits cell proliferation in all mice (Figure 4B-D; Figure S6A-B) and human (Figure 4E-G) glioma cells. Interestingly, combination treatment of AMD3100 with IR showed a greater decrease in cell viability in all GBM cells tested, compared to AMD3100 or IR alone (Figure 4B-G).

**Figure 4.**
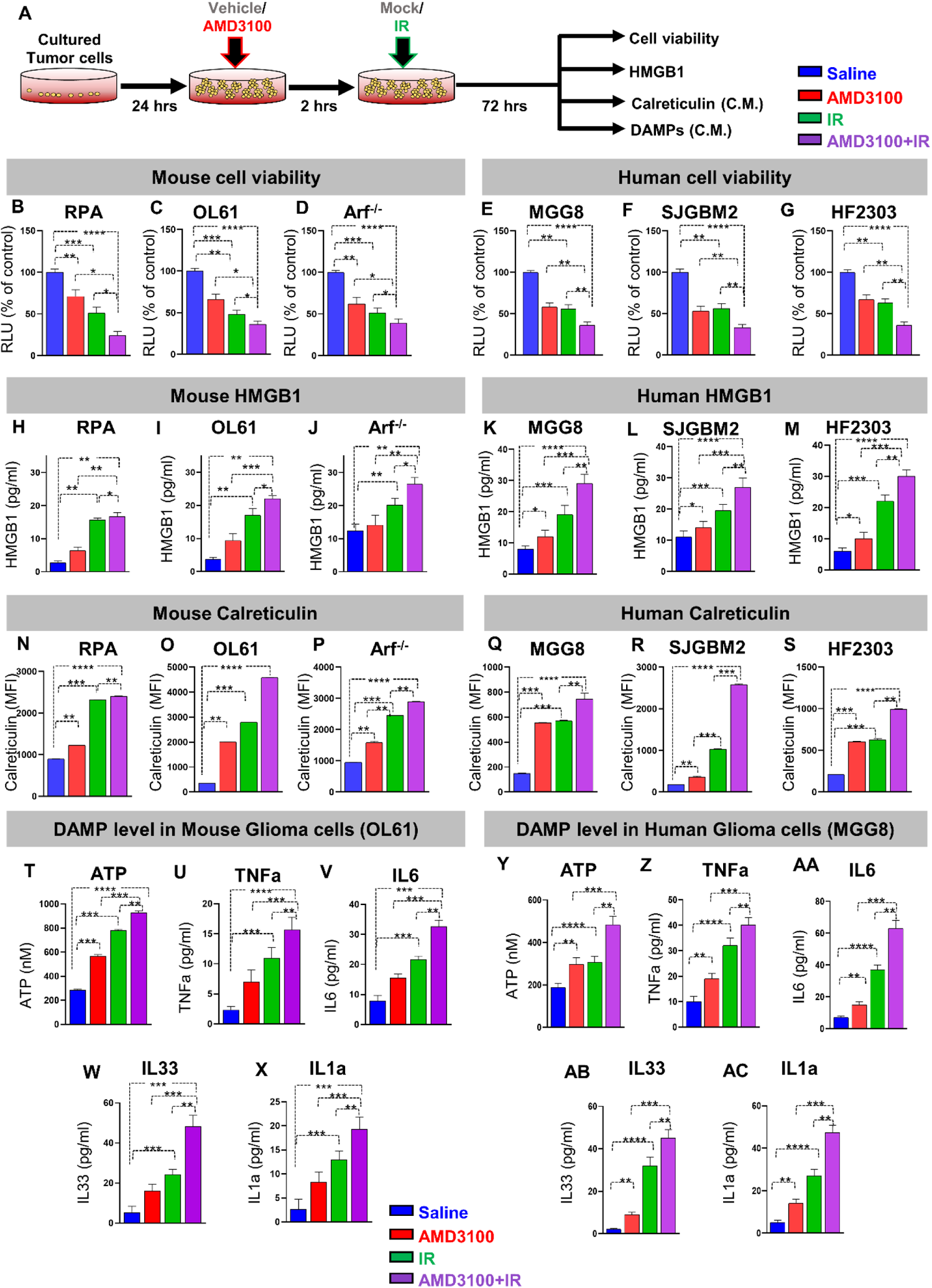
CXCR4 blockade enhances radio-sensitivity and immunogenic cell death in mouse and human glioma cells. **(A)** Schematic shows the *in vitro* application of AMD3100 and/ or radiation in mouse and human cell cultures. Mouse and patient-derived glioma cells were treated with either free-AMD3100 or in combination with radiation at their respective IC_50_ doses for 72h. All mouse and human glioma cells were pre-treated with AMD3100 2h prior to irradiation with 3Gy and 10Gy of radiation respectively. **(B-D)** Bar plot shows the % viable mouse glioma cells (RPA, OL61, or Arf^−/−^) after treatment with saline, AMD3100, IR (3Gy), or AMD3100+IR. **(E-G)** Bar plot shows the % viable human glioma cells (MGG8, SJGBM2, or HF2303) after treatment with saline, AMD3100, IR (3Gy), or AMD3100+IR. **(H-J)** Bar graphs represent levels of immunogenic cell death (ICD) marker HMGB1 in mouse glioma cells (RPA, OL61, or Arf^−/−^) after treatment with saline, AMD3100, IR (3Gy), or AMD3100+IR. **(K-M)** Bar graphs represent levels of immunogenic cell death (ICD) marker HMGB1 in human glioma cells (MGG8, SJGBM2, or HF2303) after treatment with saline, AMD3100, IR (3Gy), or AMD3100+IR. **(N-P)** Bar graphs represent levels of immunogenic cell death (ICD) marker Calreticulin in mouse glioma cells (RPA, OL61, or Arf^−/−^) after treatment with saline, AMD3100, IR (3Gy), or AMD3100+IR. **(Q-S)** Bar graphs represent levels of immunogenic cell death (ICD) marker Calreticulin in human glioma cells (MGG8, SJGBM2, or HF2303) after treatment with saline, AMD3100, IR (3Gy), or AMD3100+IR. **(T-AC)** Quantitative ELISA show the levels of DAMPs (ATP, TNFα, IL6, IL33, IL1α), as markers for ICD in the mouse glioma cells OL61 and the human glioma cells MGG8 after treatment with saline, AMD3100, IR (3Gy), or AMD3100+IR. All AMD3100 treatment were done at IC_50_ values of the corresponding cell line alone or in combination with 3Gy of IR. (Blue= Saline red= AMD3100 alone, green= IR alone, violet= AMD3100 + IR). MFI= mean fluorescence intensity. ns= non-significant, *p< 0.05 **p< 0.01, ***p< 0.0001, ****p< 0.0001; unpaired t-test. Bars represent mean ± SEM (n= 3 biological replicates).

We hypothesized that combining CXCR4 antagonist AMD3100 with radiation would induce immunogenic cell death (ICD) of glioma cells. This unique mechanism of tumor cell death results in an increased secretion of and <u>D</u>amage-<u>A</u>ssociated <u>M</u>olecular <u>P</u>atterns (DAMPs). Therefore, we measured the levels of HMGB1, Calreticulin (CRT), as well as ATP, IL1α, IL6, IL33, and TNFα in multiple GBM mouse and human cells upon treatment with saline, AMD3100, IR, or combination therapy. Mouse-GBM cells treated with AMD3100 displayed a 1.3, 4.3, and 1.6-fold (p ≤ 0.0001) increase in CRT expression relative to vehicle control group for RPA (Figure 4H), OL61 (Figure 4I), and Arf^−/−^ (Figure 4J) cells respectively. This response was further increased by approximately 2, 2.5, and 2-fold (P<0.01) with AMD3100+IR treatment for RPA, OL61, and Arf^−/−^ cells, respectively (Figure 4H-J). A similar response was observed in human-glioma cells treated with AMD3100 alone or AMD3100+IR (Figure 4K-M). We also tested the release of HMGB1 in the supernatants of mouse and human-glioma cells in response to AMD3100, IR, or AMD3100+IR treatments. Upon treatment with AMD3100, we observed a 1.5, 2, and 1.3-fold increase in HMGB1 release in the CM from RPA (Figure 4N), OL61 (Figure 4O), and Arf^−/−^ (Figure 4P) (p ≤ 0.0001) mouse glioma cells, respectively. Interestingly, combination therapy resulted in a further (~2-fold) increase in the extracellular HMGB1 release in the supernatant of all mouse glioma cells (Figure 4N-P). A similar response was observed in human-glioma cells treated with AMD3100 alone or in combination with IR (Figure 4Q-S). Next, we assessed DAMPs released from all glioma cells which received mono or dual therapy as described. Consistent with HMGB1 and Calreticulin expression, the levels of ATP, TNFα, IL6, IL33, and IL1α in all three mouse glioma cells (OL61, RPA, and Arf^−/−^) was significantly higher in the combination treatment (Figure 4T-X Figure S7A, B) and in MGG8 human-glioma cells (Figure 4Y-AC, Figure S7C, D). Taken together, these results demonstrated that AMD3100 in combination with IR enhances the radio-sensitivity and induces ICD in both mouse and human-glioma cells.

### Synthetic protein nanoparticles’ (SPNPs) design, synthesis, and characterization

AMD3100 has poor pharmacokinetic and BBB penetration. Therefore, its delivery to the GBM TME is limited ^45, 46^. To overcome this obstacle, we designed SPNPs that facilitate AMD3100 delivery to GBM. AMD3100-loaded SPNPs were prepared via electrohydrodynamic (EHD) jetting ^47–52^. In brief, a dilute solution of human serum albumin (HAS), AMD3100, and a bifunctional OEG macromer (NHS-OEG-NHS, 2 kDa) is atomized and results in the preparation of well-defined nanoparticles (Figure 5A). An aqueous HSA solution (7.5 w/v%) that also contained the OEG macromer (10 wt% relative to HSA) is mixed with 3.75 mg/ml of AMD3100. In addition, the jetting solution also contained the tumor penetrating peptide, iRGD. After jetting, the SPNPs were incubated at 37 °C to allow the OEG macromers to react with the albumin molecules resulting in water stable SPNPs. After EHD jetting, the resulting SPNPs had an average diameter of 98.5 ± 37.3 nm, as confirmed by SEM (Figure 5B, D). The PDI of these nanoparticles was 0.143, a value that is consistent with a close-to-monodisperse particle population. Furthermore, the AMD3100 SPNPs appeared to be highly spherical, as indicated by a circularity of 0.919. Once fully hydrated, the average diameter of SPNPs was 161.2 ± 46.5 nm (PDI = 0.083) based on dynamic light scattering (DLS) measurements (Figure 5C, D). Generally, AMD3100 loading had minimal influence on the particle properties. In fact, particle size, shape, and swelling behavior were very similar for AMD3100-loaded SPNPs compared to empty SPNPs (Figure 5D). The designed AMD3100 SPNPs facilitate AMD3100 entry to the GBM TME. This results in a dual anti-tumor effect by inhibiting GBM cell proliferation and decreasing infiltration of immunosuppressive CXCR4^+^ M-MDSCs (Figure 5E, F).

**Figure 5.**
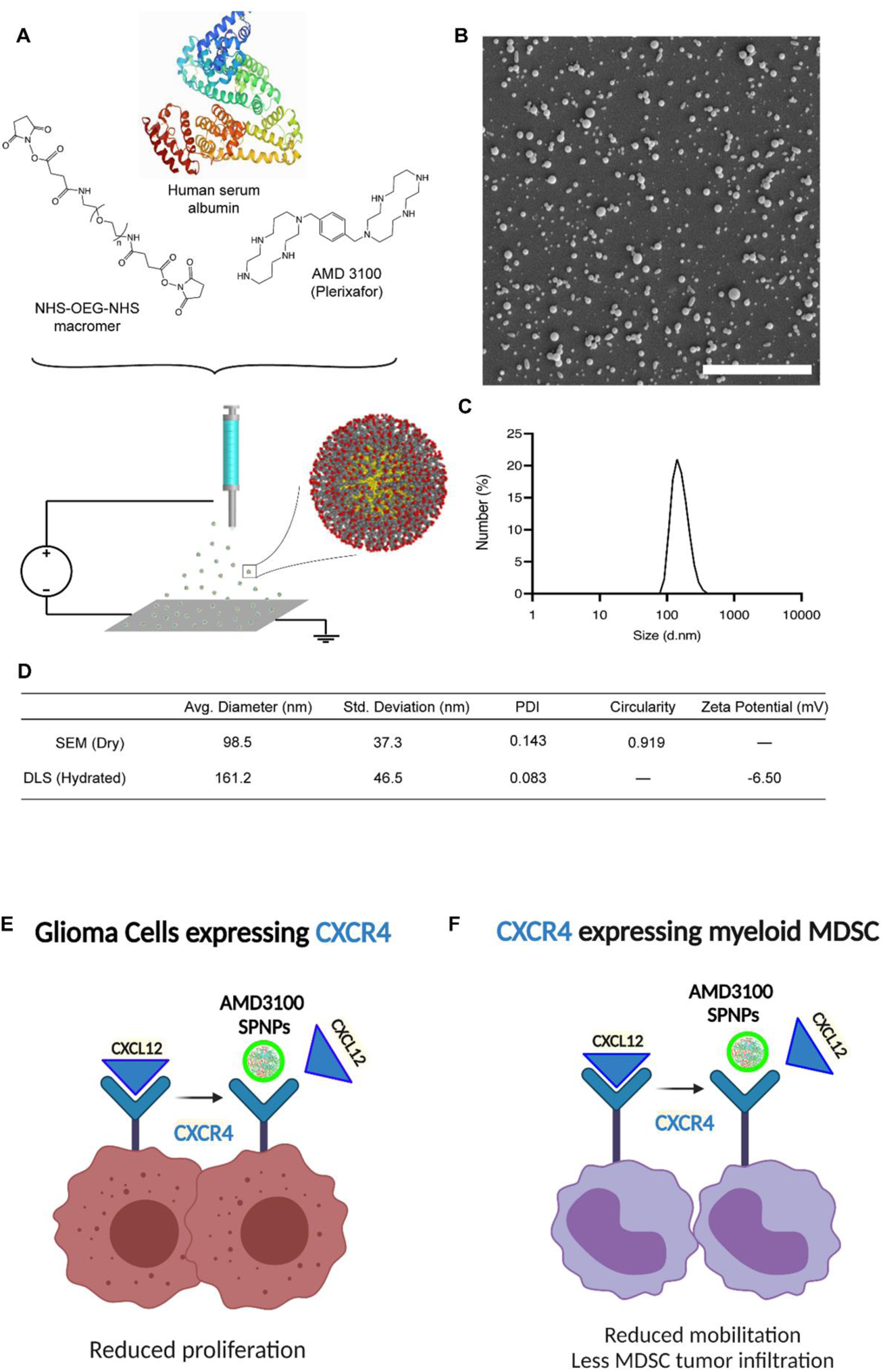
Electrohydrodynamic (EHD) co-jetting and characterization of AMD3100 SPNPs. **(A)** Schematic of the jetting process for human serum albumin SPNPs encapsulating AMD3100. **(B)** Size distribution of SPNPs in their dry state. Particle size distribution and shape characterized via Scanning Electron Microscopy (SEM) and ImageJ analysis. Average diameter, 98.5 ± 37.3 nm (PDI = 0.143). Scale bar = 3 μm. **(C)** Dynamic Light Scattering (DLS) size distribution of SPNPs in PBS. Average diameter 161.2 ± 46.5 (PDI = 0.083). **(D, E)** Schematics represent the therapeutic advantages of blocking CXCR4 in glioma cells and myeloid cells.

### AMD3100 SPNPs used in combination with radiation elicit enhanced therapeutic efficacy and anti-GBM immune response

Previous data indicated that blocking CXCR4 signaling prevents glioma progression ^10, 25, 43, 53^. To test the efficacy of AMD3100 SPNPs *in vivo*, GBM-bearing mice were treated intravenously with a total of 10 doses of AMD3100 SPNPs over the course of a three-week treatment regimen (Figure 6A). Since radiation therapy is the current standard of care for GBM, we tested the efficacy of AMD3100 SPNPs alone or in combination with two cycles of radiotherapy on the survival of tumor-bearing mice. We thus established a treatment protocol that combined the previously evaluated multi-dose of nanoparticles with a two-week focused radiotherapy regimen ^37, 40, 54^. Results showed that administration of empty SPNPs did not affect median survival compared to saline treated group (Figure 6B). Administration of AMD3100 or IR alone significantly enhanced MS of tumor-bearing mice (MS= 45 dpi for AMD3100, MS= 28dpi for IR; P<0.05, Figure 6B). However, AMD3100 SPNPs significantly enhanced survival compared to control or AMD3100 alone, underscoring the role of SPNPs in improving AMD3100 delivery to the GBM (MS= 61 dpi for AMD3100 SPNPs, MS= 45 dpi for AMD3100; P<0.05, Figure 6B). Interestingly, combining AMD3100 SPNPs with radiation resulted in a greater survival benefit with 67% of mice surviving long-term (P<0.05, Figure 6B). Long-term survivors from the combination treatment groups were rechallenged with OL-61 tumor cells in the contralateral hemisphere. These animals remained tumor-free without further treatment, whereas control mice implanted with glioma cells succumbed due to tumor burden (MS= 19 days; *P* ≤ 0.0001, Figure 6C). When sacrificed at 60 dpi after tumor rechallenge, these mice showed no evidence of microscopic intracranial tumor (Figure 6D). In contrast, H&E staining shows the presence of a fully developed tumor localized within a single hemisphere of the saline-treated mice (Figure 6D).

**Figure 6.**
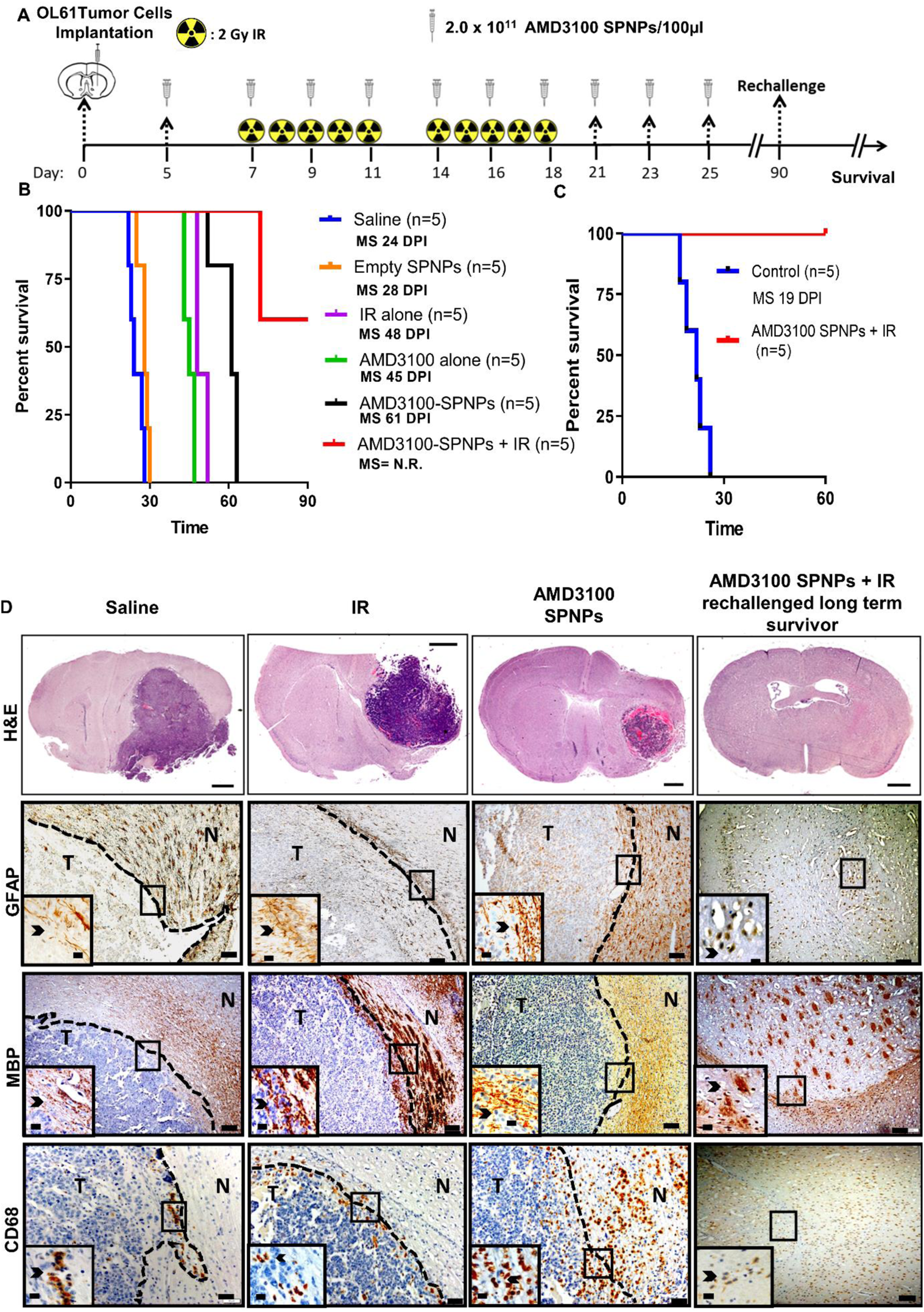
Combining AMD3100 SPNPs with IR prolong survival of GBM tumor bearing mice. **(A)** Timeline of treatment for the combined AMD3100 SPNPs+ IR survival study. **(B)** Kaplan–Meier survival curve. Significant increase in median survival is observed in all groups receiving AMD3100 alone or IR (P<0.01). Mice (3/5) treated with AMD3100 SPNPs + IR reach long-term survival timepoint (100 dpi) with no signs of residual tumor **(C)** Kaplan-Meier survival plot for re-challenged long-term survivors from AMD3100 SPNPs+IR (N=5), or control (OL61 Untreated) (N=5). Data were analyzed using the log-rank (Mantel-Cox) test. Dpi= Day post implantation. NS= Not significant. **P<0.01, ***P<0.005, One-way analysis of variance (ANOVA). **(D)** H&E staining of 5µm paraffin embedded brain sections from saline (24 dpi), IR (48 dpi), AMD3100 SPNPs alone (45 dpi) and long-term survivors from AMD3100 SPNPs + IR treatment groups (60 dpi after rechallenging with OL61 cells) (scale bar = 1mm). Paraffin embedded 5µm brain sections for each treatment groups were stained for CD68, myeline basic protein (MBP) and glial fibrillary acidic protein (GFAP). Low magnification (10X) panels show normal brain (N) and tumor (T) tissue (black scale bar = 100µm). Black arrows in the high magnification (40X) panels (black scale bar = 20µm) indicate positive staining for the areas delineated in the low-magnification panels. Representative images from a single experiment consisting of independent biological replicates are displayed.

Moreover, no apparent regions of necrosis, palisades, or hemorrhages were present in these animals 90 dpi after receiving a full course of combination therapy (Figure 6D). Myelin basic protein (MBP) and glial fibrillary acidic protein (GFAP) staining were performed to assess the integrity of myelin sheaths, an indicator for the disruption of surrounding brain architecture. No apparent changes in brain architecture were observed in mice that received the combined AMD3100 SPNPs + IR treatment when compared to the cancer-free mice in the saline-treated control group (Figure 6D, middle images). Furthermore, recognizing that significant SPNPs would be cleared via the liver ^37^, complete blood cell counts, serum biochemistry, and liver histopathological analysis were performed to examine potential off-target side effects of the combined therapeutic strategy. Systemic toxicity of AMD3100 SPNPs + IR treatment was evaluated following the treatment schedule as indicated in Figure 6A. No significant differences in the cellular composition of the blood were noted in complete blood cell counts analysis for animals receiving SPNPs, AMD3100 SPNPs, or AMD3100 SPNPs + IR treatment compared with animals in the saline treatment group (Figure S8). Moreover, there was no significant difference in biomarkers involved in the kidney (creatinine, blood urea nitrogen) and liver (aminotransferase, aspartate aminotransferase) physiology for animals receiving SPNPs, AMD3100, AMD3100 SPNPs, or AMD3100 SPNPs + IR treatment compared with animals in the saline treatment group (Figure S8), indicating that no overt adverse side-effects occurred in these tissues. Furthermore, independently conducted pathological analysis of liver tissue sections revealed no differences in the hepatocytes, stromal central, and portal areas between control saline and different treatment groups (Figure S9A-F). These results suggest that radiotherapy and AMD3100 SPNPs is a safe and effective combination for GBM tumor eradication with no cytotoxicity.

To assess the level of immune cellular infiltrates, brain tissue sections of mice from the same experimental groups were evaluated by immunohistochemistry using markers for macrophages (CD68), T-cells (CD8 and CD4), and microglia (IBA1) (Figure 6D, Figure S10). Visual inspection showed increased infiltration of CD68^+^ macrophages within the tumor and the surrounding brain parenchyma in the group that received monotherapy of either AMD3100 SPNPs or IR compared to saline treated control (Figure 6D, bottom). The long-term survivors from the AMD3100 SPNPs + IR group have a reduced number of CD68^+^ macrophages compared to the other treated groups (Figure 6D, bottom). We also observed IBA1^+^ microglia, CD8^+^ and CD4^+^ T cells within the tumor and the surrounding brain parenchyma in AMD3100 SPNPs or IR alone treated groups compared to the control group. However, there was a reduced level of IBA1^+^, CD8^+^ and CD4^+^ populations in AMD3100 SPNPs + IR long-term survivors compared to other treatment groups (Figure S10 1^st^ to 3^rd^ row). These data indicate that there was no overt inflammation in long-term survivors. Thus, our findings suggest that AMD3100 SPNPs in combination with IR is capable of eliciting an antitumor response leading to long-term survival and immunological memory.

### Combination therapy of AMD3100 SPNPs + IR primed an adaptive immune response and enhances T-cell mediated cytotoxicity

To further investigate the potential role of combination AMD3100 SPNPs and IR treatment in adaptive immunity, we examined the antigen-specific CD8 T cells within the TIME using flow cytometry. We established tumors in mice using GBM cells that harbored a surrogate tumor antigen ovalbumin (OVA) and compared the responses elicited by the various treatment formulations (Figure 7A). The OVA-based GBM model was utilized to assess the frequency of tumor antigen-specific T cells within the GBM microenvironment. Tumor-specific T cells were characterized by staining for the SIINFEKL-H2K^b^-OVA tetramer, an OVA cognate antigen within the CD8 T cell population. Results showed that the tumor-specific CD8 T cells (CD3+/CD8+/SIINFEKL-H2K^b^ tetramer) within the AMD3100 SPNPs + IR group were increased by two-fold compared to AMD3100 SPNPs alone and saline groups (P<0.01, Figure 7B). In addition, CD8 T cells infiltrating the tumor of the combination therapy have enhanced expression of the T-cell effector molecules interferon-γ (IFN-γ) and granzyme B (GzmB) (P<0.001, Figure 7C, D). Taken together with the enhanced survival observed in the combination therapy group (Figure 6B), these results suggest that a robust anti-GBM immune response was elicited by the combined AMD3100 SPNPs + IR therapy (Figure 7E). To functionally test if combining AMD3100 SPNPs with radiotherapy results in priming anti-tumor T-cell response, we co-cultured splenocytes derived from mice with the treatment formulations shown in Figure 7A, with OL61-OVA tumor cells in a ratio 20:1 (T-cell to tumor cells) *ex vivo* (Figure 7F). Results showed that splenocytes derived from combination therapy results in enhanced cytotoxicity against tumor cells compared to AMD3100 SPNPs, and saline group (P<0.05, Figure 7G). We further assessed anti-tumor T-cells elicited in combination therapy using OVA-independent OL61 model (Figure S11). Using this additional model, we found that combining AMD3100 SPNPs with radiation results in enhanced infiltration of CD3+ and CD8+ T cells (Figure S11B, C). Consistent with previous results, T-cells from combination therapy exhibited higher expression level of effector molecules (i.e. GzmB, and IFN-γ) compared to control (saline or AMD3100 SPNPs alone), but there were no differences between these effector molecules’ expression in the combination therapy and IR groups (Figure S11 D, E). These results further validated the AMD3100 SPNPs, in combination with IR, is capable of eliciting an antitumor immune response leading to long-term survival and immunological memory. Involvement of this adaptive immune response protects against secondary tumors, which is essential for successful glioma therapy that results in long-term eradication of resistant and recurrent GBM tumors.

**Figure 7.**
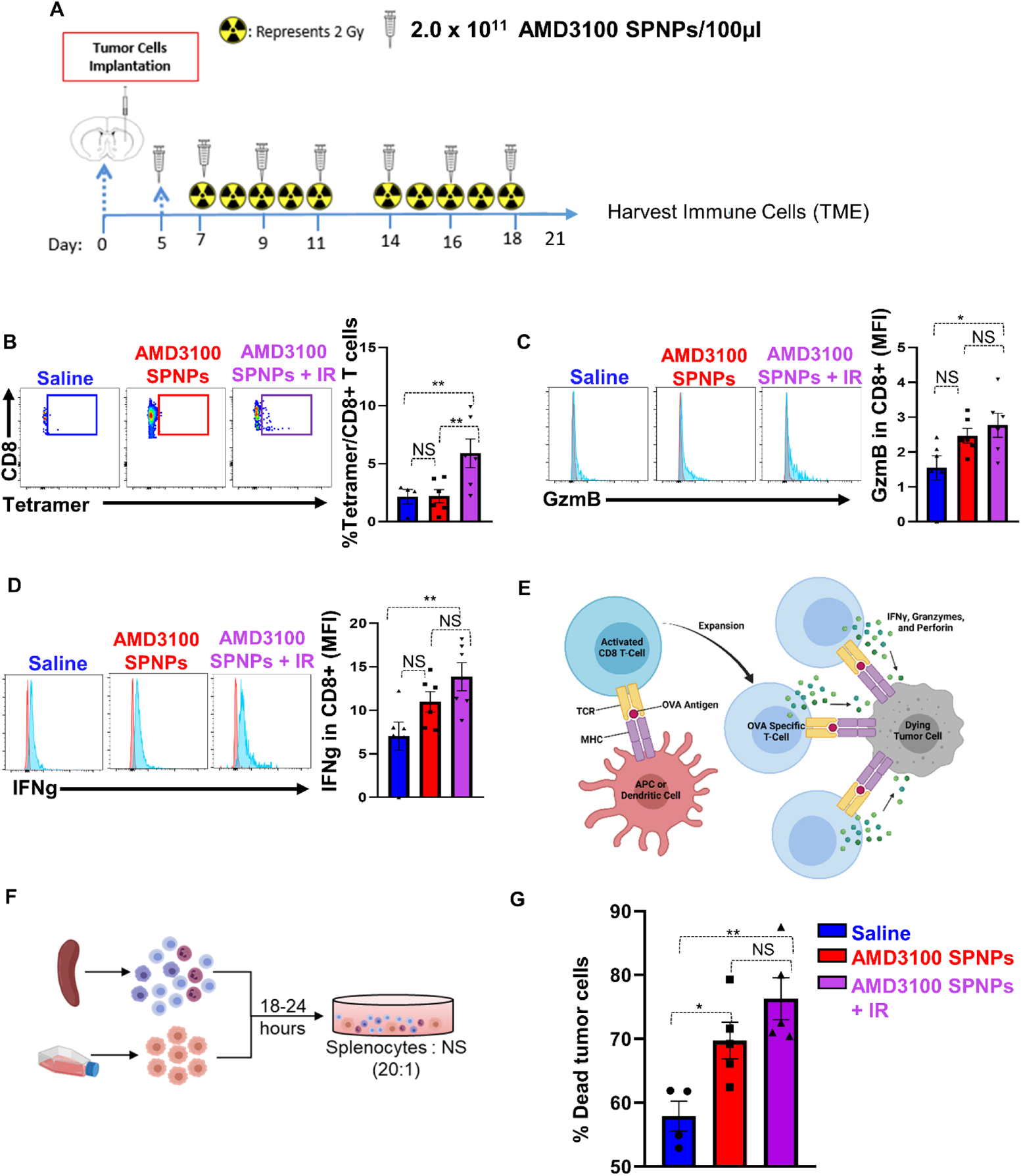
Combining AMD3100 SPNPs + IR enhances the adaptive antitumor immune response. **(A)** Experimental design represents the timeline for the combination treatment of AMD3100 SPNPs + IR to assess the efficacy of GBM-infiltrating T cell function. **(B)** Representative flow cytometry plots and analysis represents the frequency of tumor specific CD8^+^ T cells within the TME in saline, AMD3100 SPNPs, or AMD3100 SPNPs+ IR group. OL61-OVA tumors were analyzed by staining for the SIINFEKL-K^b^ tetramer. **(C, D)** Representative flow cytometry plots and analysis represent the expression of effector T cells molecules Granzyme B (GzmB) **(C)** and Interferon-g (IFN-γ) **(D)** in CD8 T cells in filtrating the TME of each group. **(E)** Schematic represents the process of priming and expansion of OVA specific CD8 T cells which target OL61-OVA cells and triggers tumor cell death. **(F)** Schematic represents the killing assay of tumor cells co-cultured with splenocytes from each treatment group. **(G)** Quantitative analysis of the percentage of tumor cells death in co-culturing condition of OL61-OVA tumor cells with Splenocytes from OL61-OVA implanted mice treated with saline, AMD3100 SPNPs, or AMD3100 SPNPs+IR. Red histogram= isotype control, blue histogram= representative sample GzmB or IFN-γ expression. *p< 0.05 **p< 0.01, ***p< 0.0001, ****p<0.0001; One way ANOVA. Bars represent mean ± SEM. (N=4-5/group).

## Discussion

The current standard of care for primary GBM involves surgical resection followed by concurrent radiation and adjuvant chemotherapy with temozolomide (TMZ) ^55, 56^. Despite therapeutic advances and intense research efforts to develop novel therapeutic modalities, the median survival of GBM patients remains poor ~18-24 months ^57, 58^. GBM patients succumb to their disease ~9-12 months post-tumor recurrence, highlighting the urgent need for developing more effective and safer anti-GBM therapies that can elicit an anti-GBM immunological memory response. We developed synthetic protein nanoparticles loaded with the CXCR4 inhibitor, i.e., AMD3100 (AMD3100 SPNPs), and demonstrated that after systemic delivery, AMD3100 SPNPs, inhibit GBM progression; when used in combination with radiotherapy they elicit GBM immunogenic cell death and an effective anti-GBM adaptive immune response. Furthermore, we showed that systemic administration of AMD3100 SPNPs resulted in a less suppressive TIME by inhibiting the infiltration of inhibitory CXCR4^+^ M-MDSCs.

Myeloid-derived suppressor cells (MDSCs) are major component of the TIME hindering effective anti-GBM immunotherapy ^59–61^. Studies showed that the number of circulating MDSCs is positively correlated with the stage of the tumor and with tumor burden, in several cancer types including glioma ^62–71^. We previously demonstrated, that depletion of immunosuppressive MDSCs significantly enhanced the efficacy of an immune-mediated gene therapy strategy ^59, 72^ which recently finished accrual as a Phase I clinical trial at our institution (ClinicalTrials.gov Identifier: NCT01811992). In the present study, we showed that CXCR4 is mainly expressed by GBM infiltrating, immunosuppressive monocytic MDSCs (M-MDSCs). Using multiple genetically engineered GBM mouse models, we showed that the frequency of CXCR4^+^ M-MDSCs infiltrating the GBM is correlated with GBM aggressiveness. This is consistent with the fact that CXCR4 expression in human patients was correlated with the glioma grade and associated with poor prognosis using both TCGA and CGGA databases.

Immunogenic cell death (ICD) represents a functionally unique response elicited by dying tumor cells which results in induction of adaptive anti-tumor immunity ^73, 74^. Our results show that blocking CXCR4 in mouse and human GBM inhibits GBM proliferation and increases the expression of HMGB1 and Calreticulin as well as the secretion of and <u>D</u>amage-<u>A</u>ssociated <u>M</u>olecular <u>P</u>atterns (DAMPS) including ATP, TNFa, IL33, and IL-6. These DAMPs directly activate macrophages and DCs by binding to cell surface receptors such as CD91, TLR2, TLR4, and the Receptor for Advanced Glycation End-products (RAGE) ^75–77^. Activation of these receptors leads to priming of both the innate and adaptive immunity which promotes the engulfing of dying cells, enhances antigen presentation, and increases the production of IL-1β ^76, 78^. All these signals are responsible for eliciting immune responses targeted to tumor cells and the establishment of antitumor immunological memory ^73, 76, 79^.

We demonstrated that CXCR4 is an important receptor for the maintenance of BBB integrity. Our data show that blocking CXCR4 on both human and mouse GBM cells, reduced GBM-mediated BBB disruption as well as decreased myeloid cells transmigration across brain microvascular endothelial cells (mBECs) *in vitro*. This was validated by intact tight junctions (Claudin-5/Zo-1) and increased paracellular resistance between mBECs monolayer upon blocking CXCR4 signaling. This illustrates that CXCL12/CXCR4 is an important signaling pathway that is hijacked by the GBM to compromise the BBB integrity, and that systemic administration of CXCR4 blocker may counteract the GBM-induced alteration in BBB permeability.

Previous studies showed that NPs can selectively target the GBM TME ^80, 81^. The SPNPs used in this study make use of a biologically compatible material albumin-based and the ability to target the GBM using iRGD, which enhances the ability of the SPNPs to penetrate the BBB and deliver small molecules to the TME. These SPNPs have been shown to facilitate delivery of therapeutics to the GBM TME, after systemic delivery, without toxicity ^37^.

CXCL12/CXCR4 promotes cancer growth and angiogenesis in multiple solid tumors. CXCR4 inhibitors are in clinical trials for the treatment of multiple solid tumors ^82, 83^. Structurally, AMD3100 is a bicyclam, composed of two cyclam rings joined by an aromatic bridge.

However, AMD3100 has poor pharmacokinetic and BBB penetration, therefore, its delivery to the GBM TME is limited ^45, 46^. Our approach using SPNPs loaded with AMD3100 facilitates delivery of AMD3100 to the brain via systemic administration with no overt side effects or cytotoxicity ^37, 84^.

Since radiation therapy is the current standard of care for GBM, we tested the efficacy of AMD3100 SPNPs in combination with radiation therapy. Our results showed that blocking CXCR4 significantly enhanced the efficacy of radiotherapy-mediated tumor cell killing both *in vitro* and *in vivo*. This was accompanied with enhanced release of DAMPs associated with ICD which elicits activation of antigen-presenting DCs to prime cytotoxic CD8^+^ T cells and leads to a robust anti–GBM response in our model. Blocking CXCR4 sensitizes GBM to ionization radiation causing enhanced GBM cell death resulting in tumor regression, long-term survival, anti-GBM immune response, and long-term immunological memory. Because of the robust anti-GBM memory response, this strategy can effectively inhibit recurrent GBM. Finally, we observed no signs of toxicity in the liver and no significant differences in the blood cells and serum chemistry relating to kidney and liver function suggesting no overt off-target side effects occurred because of the treatment.

In addition, to demonstrate the generation of tumor-antigen specific T cells in the GBM TME, we used a GBM mouse glioma model harboring the surrogate tumor antigen (Ag) ovalbumin (OVA). Using this model in which we monitored the generation of glioma Ag-specific T cells, we demonstrate an increase in the frequency of glioma–specific T cells in the TME of GBM– bearing mice treated with AMD3100 SPNPs in combination with IR. In addition, we found that the majority of T-cell infiltrating tumors with combination AMD3100 SPNPs + IR therapy express higher levels of effector molecules, i.e., GzmB and IFN-γ. These data strongly suggest that inhibiting CXCR4 in combination with IR induces an effective GBM–specific immune response.

In summary, our study brings a new mechanistic insight of the role of the CXCL12/CXCR4 signaling pathway in glioma. The use of SPNPs provides effective and minimally invasive delivery of AMD3100 to GBM bearing mice. AMD3100 SPNPs in combination with radiation elicits an effective ant-GBM response by 1) Inducing potent ICD, and 2) reprogramming the immunosuppressive microenvironment. The enhanced GBM sensitization to radiotherapy improved dendritic cell maturation promoted CD8+ T cell infiltration and distinctly boosted anti-GBM immune responses. This strategy can be harnessed to enhance immunotherapeutic efficacy in GBM patients.

## Materials and Methods

### Reagents

DMEM-F12 (11330057), DMEM (12430054), RPMI-1640 (11875119), FBS (10437028), PBS (14190250), N2 (17502048), and B27 (17504044) supplements and penicillin-streptomycin (15240062) were purchased from GIBCO, Life Technologies. Epidermal growth factor (EGF) and fibroblast growth factor (FGF) were purchased from Peprotech. Anti-mouse CD45 (147716), CD8 (100706), CD3 (100218), Gr1 (108430), CD11b (101226), Ly6G (127648), Ly6C (128028), CXCR4 (146508), GzmB (372216), and IFN-γ (505826) antibodies for flow cytometry analysis were obtained from Biolegend. SIINFEKL tetramers were obtained from MBL International (TB-5001-1). Viability dye (LIVE/DEAD™ Fixable Aqua Dead Cell Stain Kit, NC0180395) was purchased from Fisher Scientific. For immunohistochemistry, anti-mouse MBP (MAB386) and GFAP (AB5541) primary antibody was purchased from Millipore Sigma; anti-Nestin antibody (NB100-1604) was purchased from Novus biologicals; anti-mouse Iba1 (ab178846) and CD68 (ab125212) antibodies were purchased from Abcam; anti-mouse CD8a [(HS-361003(SY)] antibody was purchased from Cedarlane; anti-mouse CD4 (48274S) antibody was purchased from Cell Signaling. Alexa Fluor 647 anti-Calreticulin antibody (ab196159) was purchased from Abcam. AMD3100 was purchased from Selleck Chemical LLC (Cat# S8030). All fluorescence-conjugated antibodies used in this study are listed in Table S1.

### Synthesis of AMD3100 synthetic protein nanoparticles (AMD3100 SPNPs)

AMD3100 SPNPs were fabricated via the electrohydrodynamic (EHD) jetting process previously established in our group.^1–4^ Briefly, for all particle types, human serum albumin (HSA) was dissolved in a cosolvent mixture (80:20 v/v) of ultrapure water and ethylene glycol at a final concentration of 7.5 w/v%. A bifunctional OEG (NHS-OEG-NHS, 2kDa) and Alexa Fluor 647 labeled bovine serum albumin (BSA) were added at 10.0 and 0.5 w/w% relative to HSA to crosslink and fluorescently label the particles, respectively. Solutions prepared for the fabrication of AMD3100 SPNPs included the addition of 3.75 mg of AMD3100 per mL of jetting solution after first dissolving the drug in a small aliquot of ultrapure water. Finally, the cyclic nine amino acid tumors targeting, tissue penetrating peptide, iRGD, was integrated at 355 ng per mg of albumin by adding a stock solution of the peptide directly to the jetting solution. The complete jetting solutions were pumped through a syringe equipped with a 26-gauge blunt tip stainless steel needle at a constant flow rate of 0.2 mL h^-1^ while a constant voltage (ranging from 7.5 to 10.2 kV) was applied to form a stable Taylor cone at the needle tip. Particles were collected in grounded aluminum pans at a needle to collector distance of 15 cm and then incubated for seven days at 37°C to complete the polymerization process, yielding water-stable protein nanoparticles. AMD3100 SPNPs were then stored in dark RT conditions in their dry state for future experiments.

### Collection and purification of albumin nanoparticles

SPNPs were collected according to a standard protocol developed in our group ^1–4^. In brief, a small volume (5-10 mL) of PBS + 0.5% Tween 20, was added to the aluminum pans containing crosslinked, water-stable AMD3100 SPNPs. Gentle mechanical scraping of the pans using a plastic razor blade was used to transfer the NPs from the solid surface into the collection solution. The resulting NP suspension was briefly sonicated to disperse any aggregates and passed through a 40 μm cell straining filter. The resulting solution was centrifuged at 4,000 rpm (3,220 xg) for 4 minutes to pellet and remove any large protein aggregates. The SPNP-containing supernatant was then divided into 2 mL Eppendorf tubes and centrifuged at 15,000 rpm (21,500 xg) to pellet the NPs. Finally, the pelleted particles were combined into a single concentrated sample for use in future experiments. SPNPs were used within one week of their collection and were continually stored at 4°C in sterile PBS in the interim.

### Characterization of SPNP size, shape, and concentration

SPNPs were extensively characterized as previously described to establish reasonable tolerances against which each batch could be compared. Prior to their use in any experiments, collected particles were similarly measured to ensure they met specifications and to confirm batch-to-batch properties were consistently maintained. Physical characterization included the measurement of particle size in both their dry and hydrated state. To measure particle size, shape, and investigate their morphology, small silicon wafers were placed on the grounded aluminum collection surface and were subjected to the same seven days incubation period to complete the step-growth polymerization. These samples were imaged via Scanning Electron Microscopy (SEM) using a FEI NOVA 200 SEM/FIB instrument. SEM images were characterized using ImageJ software according to protocols previously developed in our lab ^52^. To investigate properties in their hydrated state, NPs were collected and purified as described above. The stock solution was diluted 100-fold in 0.22 μm filtered 10 mM KNO_3_ for subsequent DLS (dynamic light scattering) and zeta potential measurements. NTA (nanoparticle tracking analysis) was used to further investigate size and solution concentration. DLS, zeta potential, and NTA analyses were performed using the Malvern Nano ZSP and NanoSight NS300 instruments. AMD3100 SPNP solution concentration was further examined using the BCA (bicinchoninic acid) assay.

### AMD3100 SPNPs stability

AMD3100 SPNPs were collected and purified as described above and were studied to determine their stability under normal, short-term storage conditions. NPs from the concentrated stock solution were diluted in PBS (pH 7.4) to a particle concentration of 2.3 mg mL^-1^. The final NP solutions were stored at 4°C for a period of seven days. At regular timepoints throughout the experiment, aliquots of the stock solution were diluted 100-fold into 0.22 μm filtered 10 mM KNO_3_. Particle size distribution and zeta potential measurements were recorded and compared to determine if significant changes to either parameter were observed under normal storage conditions.

### AMD3100 SPNPs in vivo dose preparations

AMD3100 SPNPs were collected and purified as previously described prior to diluting the resulting solution to a concentration of 2.3 mg mL^-1^ in sterile PBS. Aliquots of the stock solution were diluted and standard DLS, zeta potential, and NanoSight measurements as detailed above were completed to ensure batch-to-batch consistency. Once validated, 200 μL of the stock SPNP solution was allocated per dose, per mouse for tail vein injection. Control SPNPs, encapsulating no AMD3100, were collected, purified, characterized, and dosed identically to AMD3100 SPNPs on a particle mass basis.

### Cell Line and Cell Culture Conditions

Genetically engineered mouse glioma models: RPA (*PDGFRA D842V*/ sh*TP53*-GFP/ *shATRX*), and OL61 (*shp53/PDGFB/NRAS*), OL61-OVA, were developed by the sleeping beauty model as described before ^43, 59, 85^. Arf^−/−^ wtIDH1 (*PDGFB/shP53/shATRX/Ink4a/Arf^−/−^*). Tumors were generated by injection of DF-1 (ATCC, CRL-12203) cells transfected with a combination of RCAS plasmids (*RCAS PDGFB-HA, RCAS shp53-RFP*) using the FuGENE 6 transfection kit (Roche, 11814443001) according to the manufacturer’s protocol and as previously described ^86, 87^.

Human glioma cells: HF2303 glioblastoma cells were grown in Dulbecco’s modified eagle (DMEM) media supplemented with 10% fetal bovine serum (FBS), 100 units/mL penicillin, and 0.3 mg/mL L-glutamine. SJGBM2 pediatric glioma cells were a kind gift by the Children’s Oncology Group (COG) Repository, Health Science Center, Texas Tech University. These cells were cultured in IMDM medium with L-glutamine (0.3 mg/mL) (Gibco,12440-053), 20% FBS (Gibco,10437-028), and antibiotic-antimycotic (1X) (Gibco, 15240-062) at 37 °C, 5% CO2. Cells were maintained in a humidified incubator at 95% air/5% CO2 at 37°C and passaged every 2-4 days.

### Generation of Ovalbumin Expressing Glioma Cells

Wild type OL61 glioma cells (5.0 x 10^5^) were cultured in a 6-well plate in 1mL of media (DMEM, Thermofisher Cat# 12430054; 10% FBS, FisherSci Cat# 10437028; 1x Anti-Anti, Thermofisher Cat# 15240062) with 8ug of polybrene and 100uL of 10x pLVX-cOVA-Puro (Addgene Cat#135073) lentiviral particles generated by the University of Michigan Vector Core; 1mL of media was added to the cells at 24- and 72-hours post-incubation. Once the cells became confluent, they were transferred to a larger T-25 cm^2^ flask to expand. Cells were then selected with 10ug/mL of puromycin added directly to the culture media. Cells were then passaged two times before assessing ovalbumin expression via western blot.

### Animal Strains

Six- to eight-week-old female C57BL/6 mice were purchased from Jackson Laboratory (Bar Harbor, ME) and were housed in pathogen free conditions at the University of Michigan. All experimental studies were performed in compliance with Institutional Animal Care & Use Committee (IACUC).

### Intracranial GBM Models

Tumor implantation was done as described before ^88^. Briefly, mice are anesthetized using ketamine and dexmedetomidine prior to stereotactic implantation with 50,000; 100,000 and 200,000 cells for OL61, Arf^−/−^ and RPA S8 in the right striatum respectively. The coordinates for implantation are 0.5 mm anterior and 2.0 mm lateral from the bregma and 3.0 mm ventral from the dura. Neurospheres were injected at a rate of 1 μL/min. Mice were given a combination of buprenorphine (0.1mg/kg) and carprofen (5mg/kg) for analgesia. At symptomatic stage, tumors were isolated and immune cells were characterized as described in the flow cytometry section (see below).

### Brain endothelial cells

Mouse brain microvascular endothelial cells (mBMECs) were prepared using a modified protocol already described ^89, 90^. Briefly, brains were collected from 6–8-week-old C57BL/6 mice, minced in Hanks balanced salt solution (HBSS; Invitrogen), and homogenized gently in a Dounce-type homogenizer. Myelin was removed by resuspending homogenates in an 18% dextran suspension (dextran molecular weight, 60,000 to 90,000; USB) and centrifuging. Red blood cells were removed by centrifuging isolated microvessels in a Percoll gradient (ThermoFisher) at 2,700 rpm for 11 min. The isolated microvessels were digested in HBSS solution containing 1 μg/ml collagenase/dispase (Roche), 10 U/ml DNase I (Sigma-Aldrich), and 1 μg/ml Na-*p*-tosyl-l-lysine chloromethyl ketone (TLCK) for 20 min at 37°C and precipitated with CD31-coated magnet beads (Dynabeads; ThermoFisher). These vessels were further cultured in Dulbecco’s Modified Eagle’s Medium (DMEM; Invitrogen) supplemented with 10% inactivated fetal calf serum, 2.5 μg/ml heparin (Sigma-Aldrich), 20 mM HEPES, 2 mM glutamine, and 1× antibiotic/antimycotic (all from ThermoFisher) plus endothelial cell growth supplement (BD Bioscience) and grown in 6-well plates coated with collagen type IV (BD Bioscience). This protocol typically produces primary endothelial cell cultures that are approximately 99% pure (determined by immunocytochemistry with an anti-platelet endothelial cellular adhesion molecule 1 [anti-PECAM-1] antibody; BD Bioscience).

Human brain microvascular endothelial cells (HBMEC) were obtained from iXCells Biotechnologies. Cell were plated on fibronectin coated flasks and grown in endothelial cell medium, 5% fetal bovine serum, 1% endothelial cells growth supplement and 1% penicillin/streptomycin solution in 5% CO_2_/95% air at 37°C. In all experiments, cells were used at 1^st^ or 2^nd^ passage.

### Tumor cell apoptosis assay

OL61-OVA cells were accutased, washed, counted and plated (50,000 cells) in 96 wells plate. Splenocytes from each group were washed, counted and co-cultured with OL61-OVA cells in the ratio of 1:20 (i.e50,000 OL61-OVA + 1,000,000 Sp). As a control condition OL61-OVA were cultured alone. The final volume in each well was 200 ul. The co-cultures were incubated at 37°C 5% CO_2_ overnight. At the end of the incubation period, all the wells were transferred into a V-bottom plate for staining. The staining procedure was done as described in the flow cytometry staining. Cells were washed with PBS and stained with viability dye (LIVE/DEAD™ Fixable Aqua Dead Cell Stain Kit, Cat# NC0180395). Cells were then stained with CD45 to distinguish the tumor cells from splenocytes. For AnnexinV staining, cells were washed 1x with flow buffer resuspended in 100 ul of APC-AnnexinV 1/50 diluted in AnnexinV Binding buffer. Incubated for 10 min at RT protected from light. Cells were then passed to a flow tube and 100 ul of AnnexinV binding buffer was added. Data acquisition was performed using FACS ARIA II (BD Biosciences) and analyzed using Flow Jo version 10 (Treestar).

### In vivo radiation

Ten days’ post OL61-OVA tumor cells (2 x 10^4^) implantation (bioluminescence signal= 10^6^), mice were subjected to Irradiation (IR) dose of 2 Gy for 5 days a week for two weeks for a total of 20gy of ionizing radiation. The procedure was performed as described before ^40, 54^. Briefly, mice were lightly anaesthetized with isoflurane. Mice were then placed under a copper Orthovoltage source, with the irradiation beam directed to the brain and body shielded by iron collimators. Irradiation treatment was given to mice at the University of Michigan Radiation Oncology Core.

### Cell treatment

Brain endothelial cells were cultured for 5 days to established characteristics of the brain endothelial barrier (e.g. TJ complexes and transport molecules). The mBMEC (mouse brain microvascular endothelial cells) were incubated with tumor conditioned media collected from OL61, Arf^−/−^, or RPA cells in presence or absence of AMD3100 inhibitors, while HBMEC (human brain microvascular endothelial cells) were incubated with tumor conditioned media collected from cultured HF2303 or SJGBM2 cells in presence or absence of AMD3100 inhibitors. For each condition, cells were seeded at density of 1 x 10^6^ cells into 6-well plate. Cells were then incubated with vehicle or AMD3100 (Plerixafor) at their respective IC_50_ doses for 72h in triplicate wells per condition. For irradiation, mouse and patient-derived glioma cells were irradiated with 3Gy and 10Gy, 2hrs after AMD3100 treatment, respectively. Conditioned media were then collected, centrifuged at 1000 x g for 10mins and filtered through 0.22μm syringe filter before they were incubated with mBMEC or HBMEC.

### Collection of tumor conditioned media for transmigration and cellular permeability assay

To analyze the transmigration assay and measure the cellular permeability in presence of tumor conditioned media, mouse neurospheres and human glioma cells were seeded at density of 1 x 10^6^ cells into 6-well plate. Cells were then incubated with either free-AMD3100 (Plerixafor) or in combination with radiation at their respective IC_50_ doses for 72h in triplicate wells per condition. Conditioned media were then collected, centrifuged at 1000 x g for 10mins and filtered through 0.22μm syringe filter for use.

### Cell survival analysis

Mouse OL61 (shp53/NRAS/PDGFβ), Arf^−/−^ wtIDH1(shp53/PDGFβ/Arf^−/−^) and RPA (shp53/PDGFRα/shATRX) cells and patient-derived glioblastoma cells (MGG8, HF2303 and SJ-GBM2 wtIDH1) were plated at a density of 1000 cells per well in a 96-well plate (Fisher, 12-566-00) 24h prior to treatment. Cells were then incubated with either free-AMD3100 (Plerixafor, Ontario Chemicals, Guelph, Ontario) or in combination with radiation at their respective IC_50_ doses for 72h in triplicate wells per condition. All the mouse and patient-derived glioma cells were pre-treated with AMD3100 2h prior to irradiation with 3Gy and 10Gy of radiation respectively. Cell viability was determined with CellTiter-Glo® Luminescent Cell Viability Assay (Promega, G7570) following manufacturer’s protocol. Resulting luminescence was read with the Enspire Multimodal Plate Reader (Perkin Elmer). Data was represented graphically using the GraphPad Prism software and statistical significance was determined by one-way ANOVA followed by Tukey’s test for multiple comparisons.

### DAMPs release measurement

To analyze the DAMPs in the tumor conditioned media and on tumor cells, mouse neurospheres and human glioma cells were seeded at density of 1 x 10^6^ cells into 6-well plate. Cells were then incubated with either free-AMD3100 or in combination with radiation at their respective IC_50_ doses for 72h in triplicate wells per condition. All the mouse and patient-derived glioma cells were pre-treated with AMD3100 2h prior to irradiation with 3Gy and 10Gy of radiation respectively. Release of DAMPs was assessed 72h post radiation. For assessing the level of Calreticulin (CRT), cells were collected, dissociated with Accutase and stained with anti-Calreticulin Ab (1:200) in PBS containing 2% FCS (flow buffer) for 30 min at 4^0^C. Cells were then washed 2 X with flow buffer and the fluorescence of the samples was read on a BD FACS ARIA SORP using 647 lasers. Data were analyzed with Flowjo v.10 software. Levels of HMGB1, IL33, IL6, IL1α, TNF-α in the culture supernatants was measured by ELISA according to manufacturer’s protocol (R&D) and at the Cancer Center Immunology Core, University of Michigan. Levels of ATP in the cultures’ supernatants were measured by ENLITEN ATP Assay according to manufacturer’s instructions (Promega).

### Real-time cell proliferation assay

Real-time cell analysis (RTCA) of cell proliferation was monitored using the xCELLigence DP system (Agilent). Before cell seeding, E-plates were coated with 0.5% laminin and equilibrated for 1 hour at 37°C, 5% CO2 with 100μL of respective media per well. A total of 5 × 103 cells for HF2303 and 3 × 103 cells for OL61 (shp53/NRAS/PDGFβ) and Arf^−/−^ wtIDH1(shp53/PDGFβ/Arf^-/-)^ were plated to a final volume of 200μL per well. E-plates were then equilibrated for 30 minutes.

Cells were treated with either free-AMD3100 or in combination with radiation at their respective IC_50_ doses in triplicate wells per condition before starting the experiment. All the mouse and patient-derived glioma cells were pre-treated with AMD3100 2h prior to irradiation with 3Gy and 10Gy of radiation respectively.

### Immunofluorescence and quantitative immunofluorescence analysis

For immunofluorescence staining, cell samples were preincubated in blocking solution containing 5% normal horse serum and 0.05% Triton 100X (Sigma-Aldrich) in PBS. Samples were then incubated overnight at 4°C with the following primary antibodies: anti-claudin-5 Alexa-Fluor 488 conjugated, ZO-1-Alexa Fluor 594 conjugated (both from Thermo Fisher). All samples were viewed on a confocal laser scanning microscope (Nikon A1, HSD, Japan). Quantitation of TJ-associated fragments for claudin-5 and ZO-1 were performed on 12 images obtained from three independent slides using Fiji software.

### Proximity Ligation Assay

Monolayers of mBMEC were washed, fixed and preincubated with permeabilization solution (DPBS and 0.5% Triton X-100) for 5 min following by blocking solution (DPBS++ containing 0.5% (v/v) Triton X-100 and 5% goat serum). mBMEC were then incubated with primary antibody pairs (mouse anti-ZO-1 and rabbit anti-claudin-5 antibodies) for 18 hrs. Then the mBMEC monolayers were washed with DPBS containing 5% goat serum under gentle agitation and exposed to rabbit Plus (DUO92002) and anti-mouse Minus (DUO92004) and incubated for 1 hr in a humidified incubator at 37 °C, 5% CO_2_. Protein-protein interactions were detected using the Detection Kit Red (DUO92008) according to the manufacturer’s (Sigma) instructions. Samples were incubated with ZO-1-Alexa488 antibody for labeling lateral cells border and mounted on slides using Duolink In Situ Mounting Medium with 4,6-diamidino-2-phenylindole (DUO82040) and imaged. For quantification, Fiji software was used for the automatic quantification of claudin-5-ZO-1 interaction (positive dots). Five random images were taken and analyzed to provide a representative sampling of the tissue. The number of positive dots is expressed as average Claudin-5-Zo-1 positive dots/positive cellular counts relative to the number of nuclei present in each imaging field.

### Permeability assays

The permeability of brain endothelial cell monolayers to cadaverin-569 (1kDa), Dextran (Cascade blue 3kDa, Thermo Fisher, D7132), Dextran Texas Red (10kDa) (Thermo Fisher, D1828), and FITC-inulin (5 kDa, Sigma Aldrich, I3661), were measured as described in Kazakoff et al., and modified in our laboratory ^91, 92^. Briefly, hBMEC or mBMEC were plated on a Transwell Dual chamber system at a density of 1×10^5^. Monolayer permeability (transendothelial electrical resistance) was measured daily from 1–7 days. Permeability experiments were initiated by adding a cocktail of tracers (1 μg/ml), in phenol red free DMEM (Thermo Fisher) in the apical chamber. Media was sampled after 30 minutes from the receiving chamber. Fluorescence intensity was determined by fluorescent reader (Tecam) and concentration determined from a standard curve. The permeability (Papp; cm/min) of the monolayer for each time points (T) was calculated using the following equation: P=(C(B)T−C(B))×V(B)×2 /(C(A)+C(A)T)×A; where C(B) and C (B)_T_ are, respectively, the concentrations of tracer in the basal chamber at the start and at the end of the time interval of 30 min (in μg/ml), and V(B) is the volume of the basal chamber (in ml). C(A) and C(A)_T_ are, respectively, the tracer concentrations in the apical (donor) chamber at the start and at the end of the time interval of 30 min (in μg/ml) and A is the area of the filter (cm^2^).

### Complete Blood Serum Biochemistry

Blood from OL61 OVA glioma bearing mice was taken from the submandibular vein and transferred to serum separation tubes (Biotang). Samples in the serum separation tubes were left at room temperature for 60mins to allow for blood coagulation before centrifugation at 2000 rpm (400 x g). Complete serum chemistry for each sample was determined by in vivo animal core at the University of Michigan.

### Immunohistochemistry

For neuropathological assessment, brains fixed in 4% paraformaldehyde (PFA) were embedded in paraffin and sectioned 5µm thick using a microtome system (Leica RM2165). Immunohistochemistry was performed on brain sections by permeabilizing them with TBS-0.5% Triton-X (TBS-Tx) for 20 min. This was followed by antigen retrieval at 96 °C with 10 mM sodium citrate (pH 6) for an additional 20 min. Then, the sections were cooled at room temperature (RT) and washed 5 times with TBS-Tx (5 min per wash) and blocked with 10% goat serum in TBS-Tx for 1 h at RT. Brain sections were incubated with primary antibody CD4 (Cell Signaling, 48274S, 1:1000), GFAP (Millipore Sigma, AB5541, 1:200), MBP (Millipore Sigma, MAB386, 1:200), IBA1 (Abcam, ab178846, 1:1000), CD8 (Cedarlane, [(HS-361003(SY)], 1:100),CD68 (Abcam, ab125212 1:1000) or Nestin (Novus Biologicals, (NB100-1604), 1:1000) diluted in 1% goat serum TBS-Tx overnight at RT. On the next day sections were washed with TBS-Tx 5 times. Brain sections labeled with CD4, GFAP and IBA1 were incubated with biotinylated secondary antibody, while brain sections labeled with CD8, CD68, MBP and Nestin were incubated with HRP secondary antibody, which were diluted 1:1000 in 1% goat serum TBS-Tx in the dark for 4h. HRP and biotin-labeled sections were subjected to 3, 3′-diaminobenzidine (DAB) (Biocare Medical) with nickel sulfate precipitation. The reaction was quenched with 10% sodium azide; sections were washed 3 times in 0.1 M sodium acetate followed by dehydration in xylene, and coverslipped with DePeX Mounting Medium (Electron Microscopy Sciences). Images were obtained using brightfield microscopy (Olympus BX53) at 10X and 40X magnification.

For tumor size quantification, paraffin embedded 5µm thick brain sections from each experimental groups were stained with H&E as described previously^93^. Sections comprising of tumor or from rechallenged brains (approximately 10-12 sections per mouse) were imaged using the brightfield (Olympus BX53) setting and tumor size was quantified using ImageJ’s Otsu threshold to determine the tumor size in pixels.

For histological assessment, livers were embedded in paraffin, sectioned 5µm thick using the microtome system and H&E stained as described by us previously^93^. Brightfield images were obtained using Olympus MA BX53 microscope.

### Flow Cytometry

For flow cytometry analysis, cells within the TME, spleen and blood from tumor bearing mice were processed as described before ^94, 95^. Tumor mass within the brain, spleen, and blood were carefully collected and homogenized using Tenbroeck (Corning) homogenizer in DMEM media containing 10% FBS. Tumor infiltrating immune cells was enriched with 30%70% Percoll (GE Lifesciences) density gradient. Cells were resuspended in PBS containing 2% FBS and non-specific antibody binding was blocked with FC block (CD16/CD32). PMN-MDSCs were labelled as CD45^high^/CD11b^+^/Ly6G^+^/Ly6C^low^, whereas M-MDSCs were gated as CD45^high^/ CD11b^+^/Ly6G^-^/Ly6C^high^. Tumor-specific T-cells were labeled with CD45, CD3, CD8 and SIINFEKL-H2Kb-tetramer. Activated T cells were labeled with CD45, CD3, CD8 and GzmB antibodies. Intracellular Granzyme B and IFN-γ were stained using BD intracellular staining kit using the manufacturer’s instructions. All stains were carried out for 30min at 4°C with 3X flow buffer washes between live/dead staining, blocking, surface staining, cell fixation, intracellular staining and data acquisition. Flow data has been measured with FACSAria flow cytometer (BD Bioscience) and analyzed using Flow Jo version 10 (Treestar).

### Western blots

Ovalbumin expressing OL61 glioma cells (1.0 x 10^6^ cells) were harvested, and total protein extracts were prepared in RIPA lysis buffer with 1X of protease/phosphatase inhibitor cocktail. 10 µg of protein extract determined by bicinchoninic acid assay (BCA kit, Thermofisher Cat# PI23227), were separated by 4-12% SDS-PAGE and transferred to nitrocellulose membranes. The membrane was probed with specific antibodies described in the figure (Vinculin, Thermofisher Cat# PI700062; Ovalbumin, Abcam Cat# ab181688), then followed by anti-Rabbit HRP secondary (Agilent, Cat# P044801-2) at 1:4000 diluted with 5% bovine serum albumin solution. Enhanced chemiluminescence reagents were used to detect the signals. WB quantification was performed using ImageJ, and the reported data are from three individual biological replicates.

### TCGA and CGGA survival analysis

Glioma TCGA dataset were downloaded from Gliovis portal http://gliovis.bioinfo.cnio.es/ glioma patients expressing high vs low *CXCR4* were stratified based on the median cut off expression value. All CGGA data were downloaded from the Chinese Glioma Genome Atlas http://www.cgga.org.cn/ and stratified in a way similar to TCGA data.

### Quantitative ELISA

Conditioned media from mouse and human glioma cells were harvested after culturing of 2 x 10^5^ cells/ 1mL for 48 hours in appropriate culture media. Quantitation of CXCL12 was determined by ELISA (Duosets, R&D Systems, Minneapolis, MN) using the manufacturer’s suggested protocol with few modifications. Briefly, diluted coating Ab was added to ELISA microplates (Greiner Bio-One, Monroe, NC) and incubated overnight. Assay plates were then washed, blocked, and samples and standards added and incubated overnight at 4° C. Diluted secondary Ab was added after washing and incubated for 3 hours at room temperature, followed by washing and HRP incubation for 90 minutes. Following a final series of washes, plates were developed with TMBX substrate (Surmodics, Eden Prairie, MN) and stopped by the addition of an equal volume of 0.4% NaF. Absorbances were obtained at 620 nm and sample concentrations were determined by comparison to the CXCL12 standards using a 4-parameter curve fit (Synergy HT & Gen5 Software, BioTek Instruments, Winooski, VT). Serum level of CXCL12 was assessed using undiluted serum isolated from the blood of tumor-bearing mice or stage IV glioma patients.

### Statistical Analysis

Sample sizes were chosen based on preliminary data from pilot experiments and previously published results in the literature. All animal studies were performed after randomization. Data were analyzed by one- or two-way analysis of variance (ANOVA), followed by Tukey’s multiple comparisons post-test or log-rank (Mantel-Cox) test with Prism 8.1 (GraphPad Software). Data were normally distributed and variance between groups was similar. P values less than 0.05 were considered statistically significant. All values are reported as means ± SD with the indicated sample size. No samples were excluded from the analysis.

### Associated Content

#### Supporting Information

The Supporting Information is available free of charge on the [ACS Publications website](http://pubs.acs.org)

➢ Table S1: Fluorescence-conjugated antibodies used in the study, Figure S1: Dose-response curves for mouse and human glioma cells treated with AMD3100, Figure S2: AMD3100 preserves the trans-endothelial impermeability of the brain endothelial cells (BEC), Figure S3: AMD3100 protects the tight junction molecules (Zo-1, and Claudin-5) between brain endothelial cells (BEC) from tumor-mediated loss of tight junctions, Figure S4: CXCR4 signaling causes derangement in tight junction molecules, Figure S5: CXCR4 signaling disrupts human brain endothelial cell (hBEC) barrier, Figure S6: Real-time cell proliferation analysis of mouse neurospheres and human glioma cells following AMD3100 + IR treatment, Figure S7: CXCR4 antagonist, AMD3100 in combination with radiation increases the release of DAMPs from both human and mouse glioma cells, Figure S8: Mouse serum biochemical analysis following intravenous AMD3100 SPNP in combination with IR treatment, Figure S9: Histopathological assessment of livers from tumor bearing mice treated with AMD3100 SPNPs + IR, Figure S10: AMD3100 SPNPs treatment in combination with radiation enhances immune infiltration of glioma bearing mice, and Figure S11: combining AMD3100 SPNPs + IR enhances the adaptive antitumor immune response.

### Statistical Analysis

Sample sizes were chosen based on preliminary data from pilot experiments and previously published results in the literature. All animal studies were performed after randomization. Data were analyzed by one- or two-way analysis of variance (ANOVA), followed by Tukey’s multiple comparisons post-test or log-rank (Mantel-Cox) test with Prism 8.0 (GraphPad Software). Data were normally distributed and variance between groups was similar. P values less than 0.05 were considered statistically significant. Unless otherwise indicated, all values are reported as means ± SEM with the indicated sample size. No samples were excluded from the analysis.

### Associated Content

#### Supporting Information

The Supporting Information is available free of charge on the [ACS Publications website](http://pubs.acs.org)

## Author Contributions

M.S.A., K.B., A.A.M., A.T., R.T., B.L.M., S.M.T., G.M-R, P.K. A.C., J.A.J., A.A.A., S.C., A. C., H.A., J.V.G., M.B.E., M.R.O. and S.Y.M., performed experiments and analyzed the data. M.S.A., K.B., A.A.M., E.RL., A.A-Z, J.J.M., P.R.L., M.G.C. designed the research and contributed to writing the manuscript. M.S.A., K.B., A.A.M., M.L.V., J.V.G., M.B., and M.G.C. designed and organized the figures. M.S.A., K.B., A.A.M., P.K., B.L.M., and M.G.C. proof-read and revised the manuscript.

## Acknowledgments

This work was supported by “National Institutes of Health/National Institute of Neurological Disorders & Stroke (NIH/NINDS) Grants [R37-NS094804, R01-NS105556, R21-NS107894 and Rogel Cancer Center Scholar Award to M.G.C.]; NIH/NINDS Grants [R01-NS076991, R01-NS082311, and R01-NS096756 to P.R.L.]; the Department of Neurosurgery; the Pediatric Brain Tumor Foundation, Leah’s Happy Hearts Foundation, Ian’s Friends Foundation (IFF), Chad Tough Foundation, and Smiles for Sophie Forever Foundation to [M.G.C. and P.R.L]. NIH/NCI T32-CA009676 Post-Doctoral Fellowship to M.S.A. NIH/NCI F31CA247104 to AAA, and NIH/NCI F31CA247104 to JAJ

## Supporting Information

**Supplementary Table S1.** Fluorescence-conjugated antibodies used in the study

| **Antibody** | **Manufacturer** | **Cat #** |
| --- | --- | --- |
| **PE/Cyanine 7 anti-mouse IFN-γ** | BioLegend | 505826 |
| **PE/Dazzle 594 anti-human/mouse Granzyme B Recombinant** | BioLegend | 372216 |
| **PerCP/Cy5.5 anti-mouse/human CD11b** | BioLegend | 101228 |
| **APC anti-mouse CD184 (CXCR4)** | BioLegend | 146508 |
| **PE/Cy7 anti-mouse Ly-6C** | BioLegend | 128018 |
| **PE/Dazzle 594 anti-mouse Ly6G** | BioLegend | 127648 |
| **Pacific Blue anti-mouse Ly6G/Ly6C (Gr-1)** | BioLegend | 108430 |
| **FITC Rat Anti-Mouse CD184 (CXCR4)** | BD Biosciences | 551967 |
| **APC/Cyanine7 anti-mouse/human CD11b** | BioLegend | 101226 |
| **APC anti-mouse CD45** | BioLegend | 103111 |
| **PerCP anti-mouse Ly-6G** | BioLegend | 127654 |
| **iTag MHC Tetramer H-2Kb OVA SIINFEKL-PE** | MBL | TB-5001-1 |
| **CD8 alpha Monoclonal Antibody (KT15), FITC** | Invitrogen | MA5-16760 |
| **Pacific Blue anti-mouse CD45** | BioLegend | 103126 |
| **PerCP/Cyanine5.5 anti-mouse TCR B chain** | BioLegend | 109228 |

**Figure S1.**
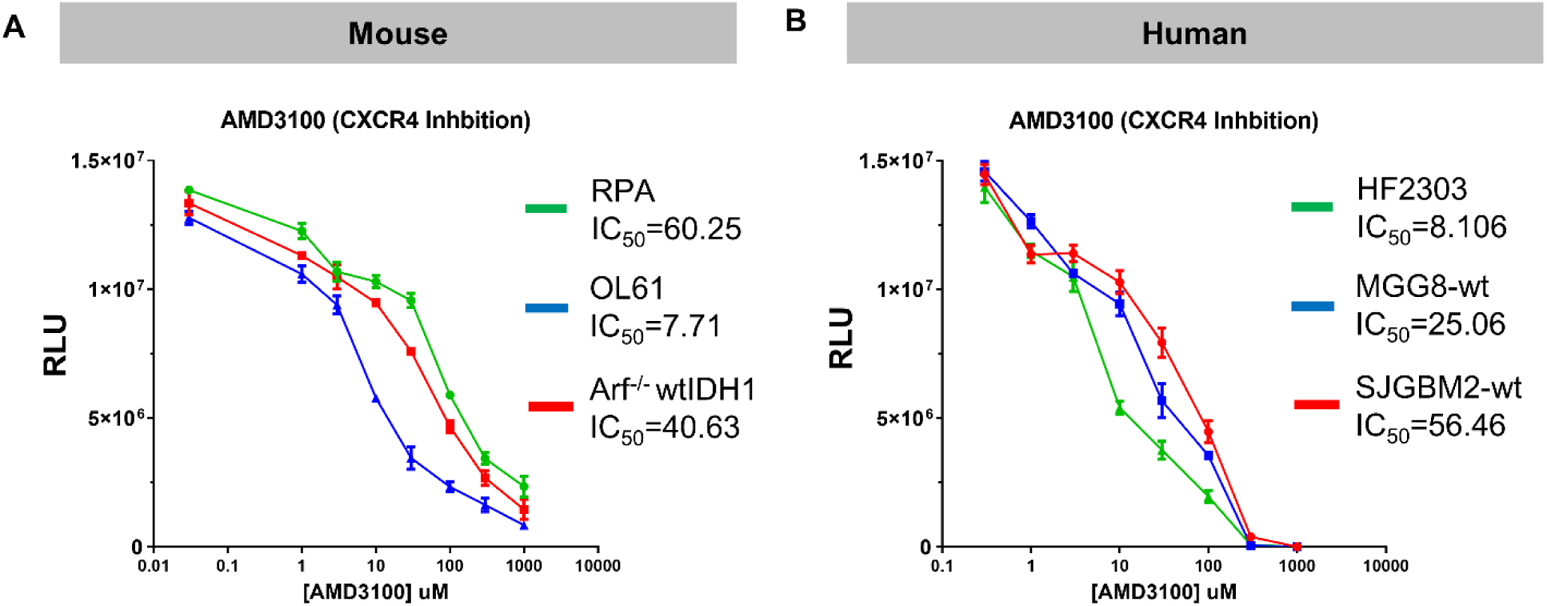
Dose-response curves for mouse and human glioma cells treated with AMD3100. Glioma mouse cells **(A)** or patient-derived human glioma cells **(B)** were incubated for 72 hrs with AMD3100 at the indicated doses. Cell viability was assessed using Promega Cell Titer Glo Assay. GraphPad Prism was used to determine IC_50_ values of the compound’s cytotoxicity, shown in μM.

**Figure S2.**
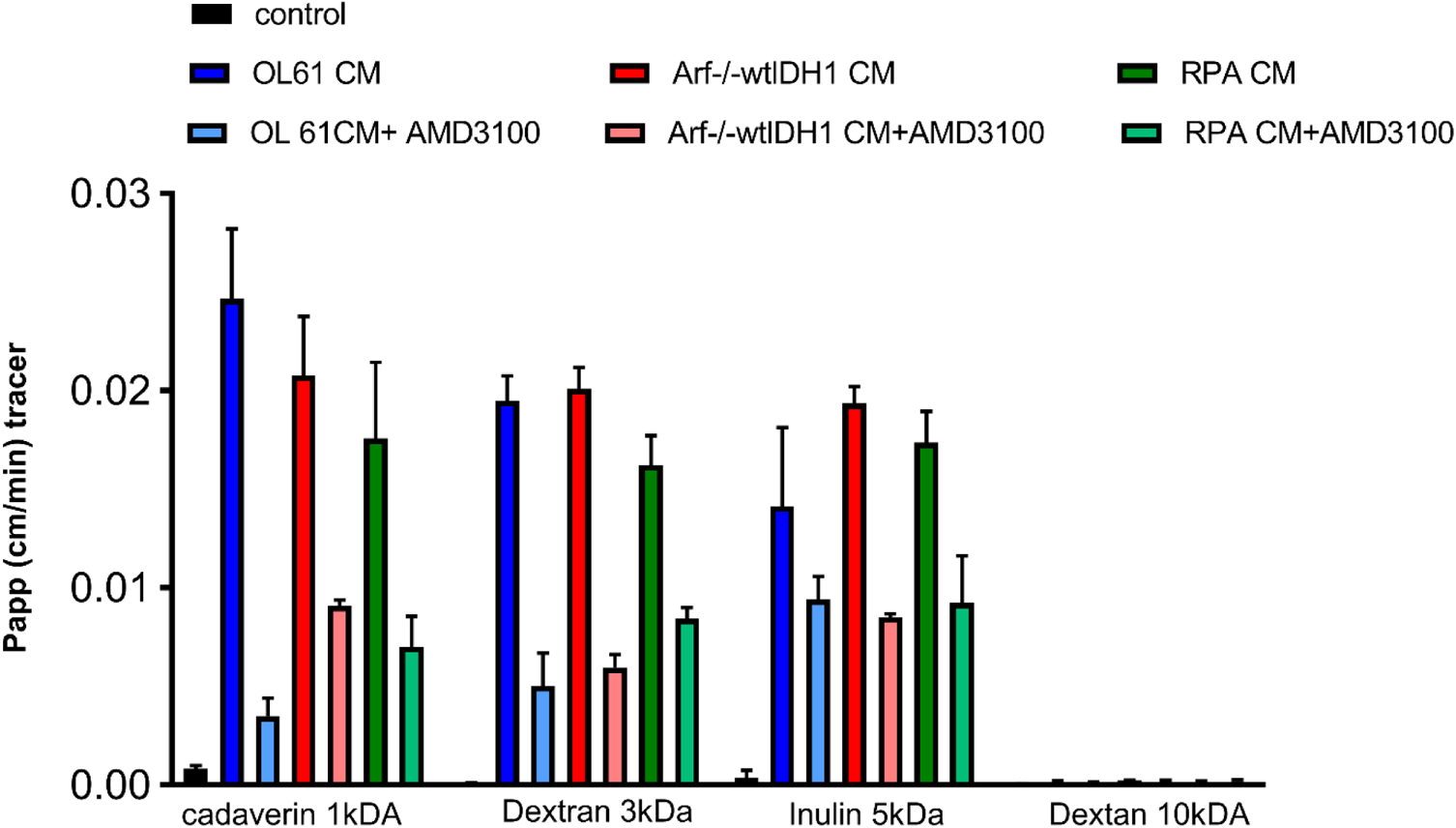
AMD3100 preserves the trans-endothelial impermeability of the brain endothelial cells (BEC). Summary data for permeability coefficient (Papp) for all tested tracers Cadaverin (1kDa), Dextran (3kDa), Inulin (5kDa), and Dextran (10kDa). The brain endothelial cells barrier showed permeability for tracer Cadaverine (1kDa), Dextran (3kDa), and Inulin (5kDa). However, the brain endothelial monolayers were not permeable for tracer Dextran 10kDa, indicating rather leakage (small molecular weight tracer permeability) than barrier breakdown (high molecular weight permeability) Data are shown as means ± SD. *n* = 3

**Figure S3.**
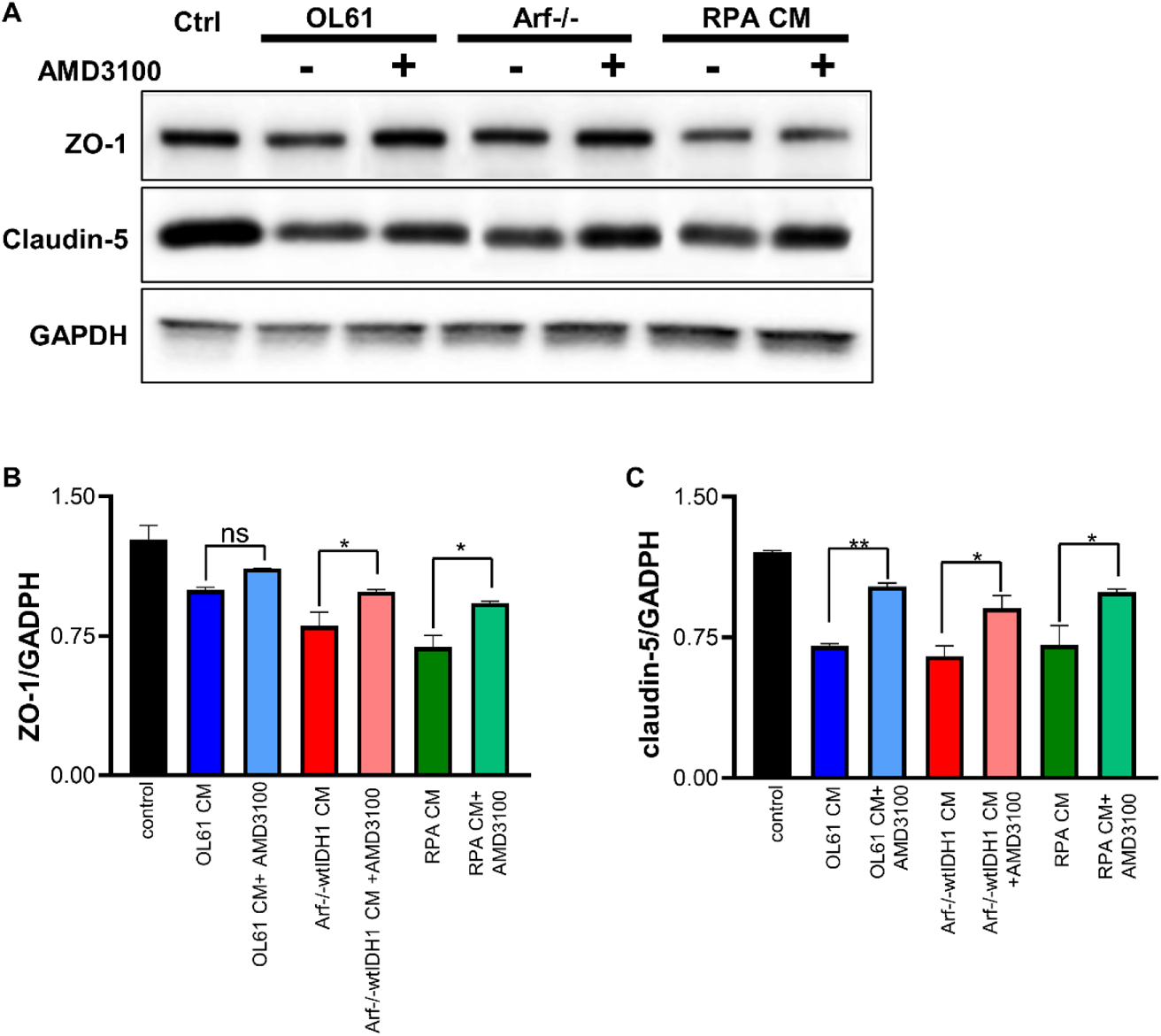
AMD3100 protects the tight junction molecules (ZO-1, and Claudin-5) between brain endothelial cells (BEC) from tumor-mediated loss of tight junctions. **(A)** Western blot analysis of claudin-5 and ZO-1 expression in control, OL61, OL61+AMD3100, Arf^−/−^wtIDH1, Arf^−/−^wtIDH+AMD3100, and RPA and RPA+AMD3100 experimental groups. Representative images of Western blot of two independent experiments are shown. **(B, C)** Bar graph shows semiquantitative densitometric analysis of ZO-1 and claudin-5. Data are shown as means ± SD, n=2 independent experiment.

**Figure S4.**
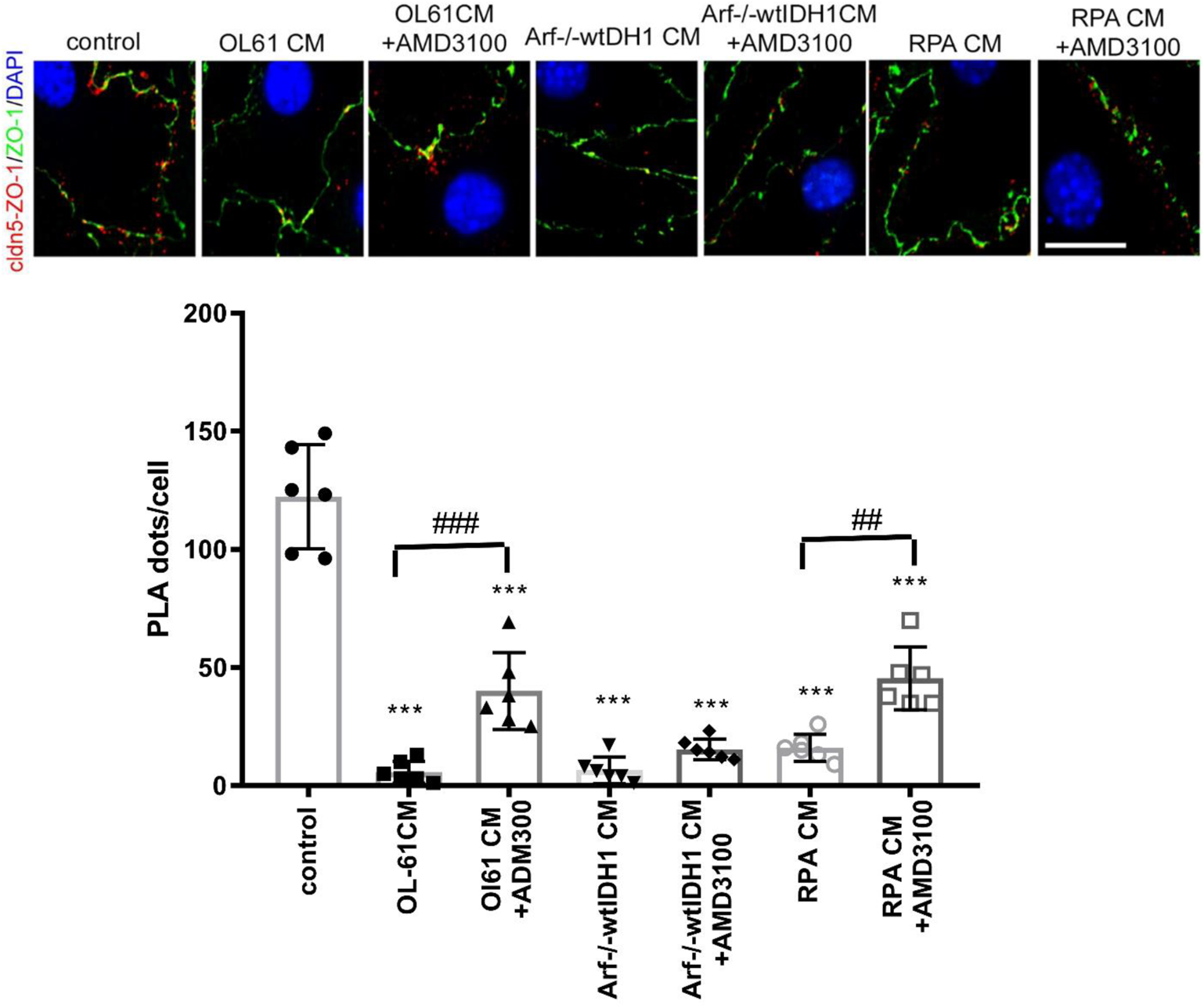
CXCR4 signaling causes derangement in tight junction molecules. **(A)** Claudin-5/ZO-1 interaction analyzed by proximity ligation assay (PLA) in control (non-exposed cells), to OL61, OL61+AMD3100, Arf^−/−^wtIDH,1, Arf^−/−^wtIDH+AMD3100, and RPA and RPA+AMD3100 for 24 hrs. mBEC monolayers were PLA labeled using claudin-5 and ZO-1 antibodies. Claudin-5 and ZO-1 interaction (cldn5-ZO-1, red dots) were mostly present on cell borders (labeled with ZO-1, green) in control cells. This reflects the claudin-5 and ZO-1 interaction in the Tj complex. OL61, Arf^−/−^wtIDH, and RPA condition media cause the barrier impairment and decreased presence of interaction site of claudin-5 and ZO-1. Condition media with AMD3100 has a protective effect indicating more interaction and increase stability of the Tj complex. Scale bar 50 μm. **(B)** The PLA signals were counted with the Image J software and the average number of spots per cell is presented in the graph. Data are shown as means ± SD, n=3 independent experiments (slides). Data are shown as means ± SD. *n* = 30 cells; \*\*\**p* = 0.0001 compared with control, ##p>0.001, ### p> 0.0001 compared cell with or without treatment with AM3100.

**Figure S5.**
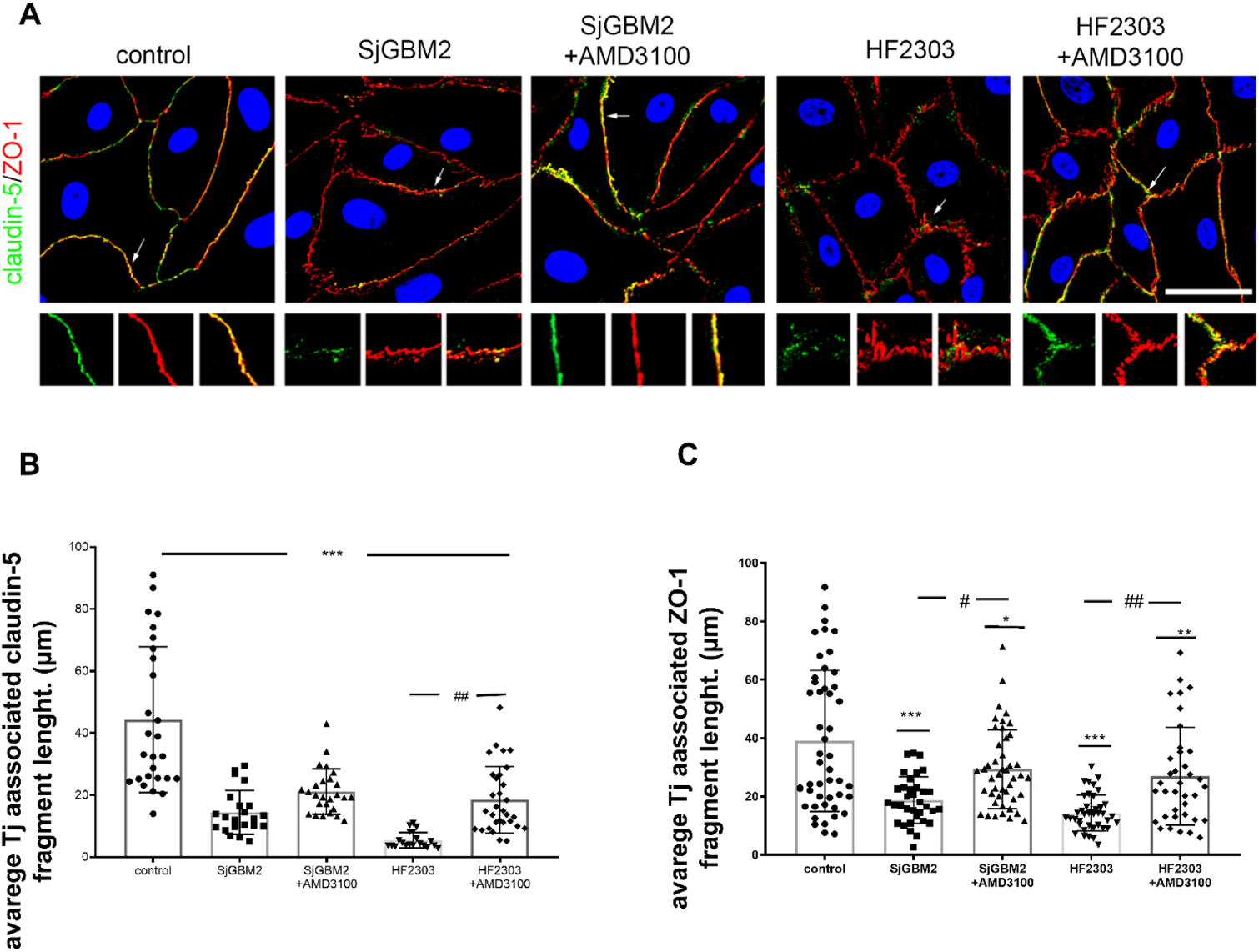
CXCR4 signaling disrupts human brain endothelial cell (hBEC) barrier. **(A)** Immunofluorescence staining for tight junction (Tj) proteins claudin-5 and ZO-1 in control and cells exposed to OL61, OL61+AMD3100, Arf^−/−^ wtIDH, Arf^−/−^ wtIDH+AMD3100, and RPA and RPA+AMD3100 for 24 hrs. Arrow and magnified images indicate pattern and colocalization of ZO-1 and claudin-5 on the cell border. Scale bar 50μm. Quantitation of the average TJ-associated **(B)** claudin-5 and **(C)** ZO-1 fragment length in claudin-5/ZO-1 costained immunofluorescent images in control and cells exposed to OL61, OL61+AMD3100, Arf^−/−^ wtIDH, Arf^−/−^ wtIDH+AMD3100, and RPA and RPA+AMD3100 for 24 hrs. Data are shown as means ± SD. *N* = 5; ***p>0.0001 and **p>0.001 comparing to control. ###p>0.0001 comparing experimental groups with and without inhibitor AMD3100.

**Figure S6.**
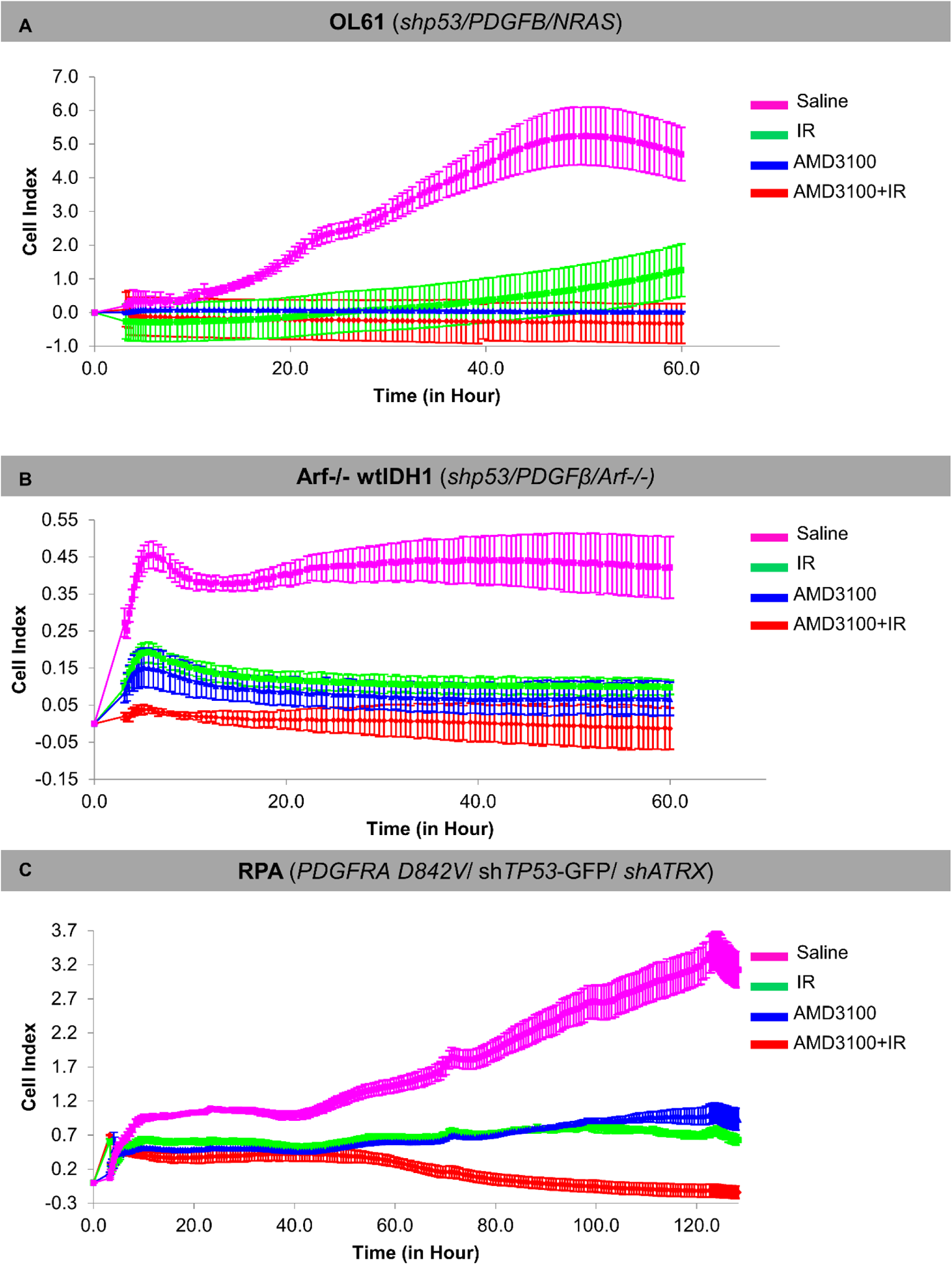
Real-time cell proliferation analysis of mouse neurospheres and human glioma cells following AMD3100 + IR treatment. Impact of AMD3100 alone or in combination with IR on cellular proliferation were analyzed through Real-time cell analysis (RTCA) xCelligence proliferation assay on (A) OL61 (shp53/NRAS/PDGFβ), (B) Arf^−/−^wtIDH1(shp53/PDGFβ/ Arf^−/−^), **(B)** HF2303 cells with AMD3100 (Blue line), IR (Green line) and AMD3100 + IR (Red line) in comparison to non-treated non-irradiated control (Pink line). OL61 (shp53/NRAS/PDGFβ) and Arf^−/−^ wtIDH1(shp53/PDGFβ/ Arf^−/−^) cells were irradiated with 3Gy and HF2303 were irradiated with 10Gy of radiation. Cells were treated with AMD3100 at their respective IC_50_ values. Cells were incubated within chambers for the indicated time periods and each time point represents the mean of technical quadruplicates. Data represents the mean of 3 independent biological replicates. Statistical analysis was done through two-way ANOVA. ***P<0.001 level of significance.

**Figure S7.**
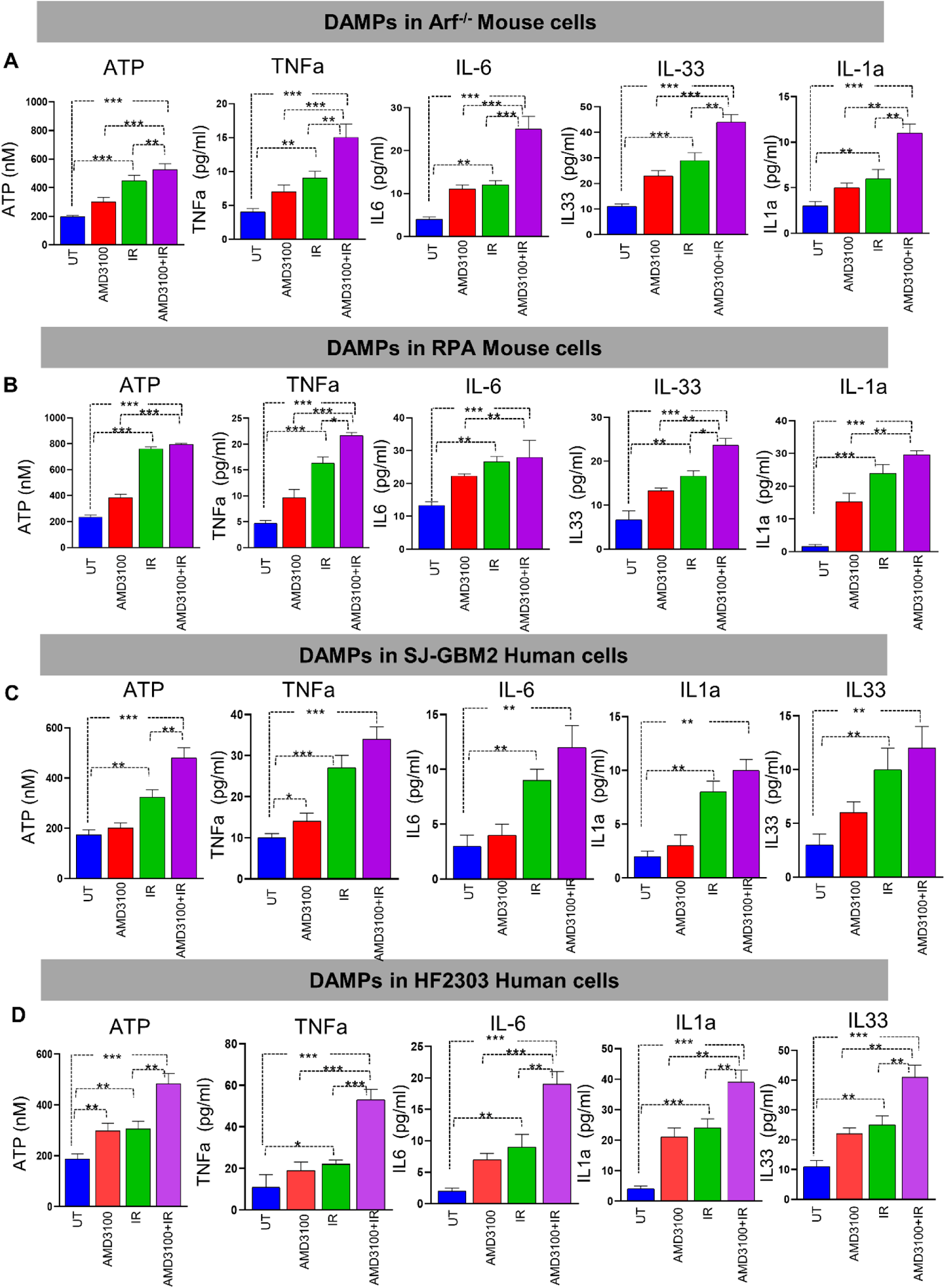
CXCR4 antagonist, AMD3100 in combination with radiation increases the release of DAMPs from both human and mouse glioma cells. Levels of DAMPs released as ICD markers i.e., ATP, TNFα, IL6, IL33, IL1α from mouse glioma cells (A) Arf^−/−^ wtIDH1, (B) RPA wtIDH1 and human glioma cells (C) SJ-GBM2 wtIDH1, (D) HF2303 wtIDH1 were determined through ELISA following AMD3100 (at IC_50_ doses) in combination with IR (3Gy and 10Gy of radiation respectively) treatment. Representative bar diagrams display each marker’s expression levels (bar diagrams: red= AMD3100 alone, green= IR alone, violet= AMD3100 + IR) compared to non-treated, non-irradiated controls (blue= untreated). ns= non-significant, *p< 0.05 **p< 0.01, ***p< 0.0001, ****p< 0.0001; unpaired t-test. Bars represent mean ± SEM (n= 3 biological replicates).

**Figure S8.**
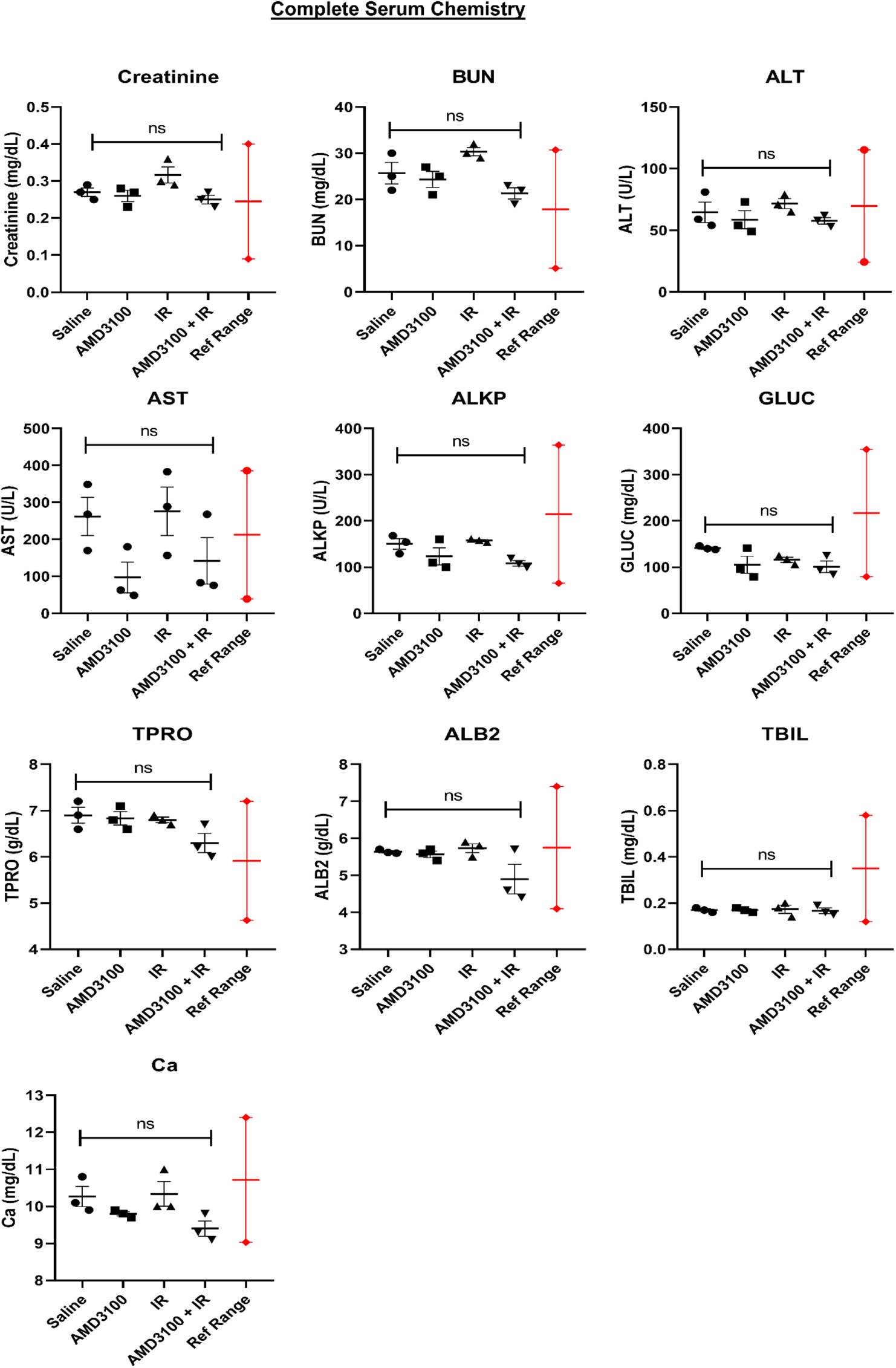
Mouse serum biochemical analysis following intravenous AMD3100 SPNP in combination with IR treatment. OL61 OVA tumor bearing mice treated with CXCR4 antagonist AMD3100 SPNPs in combination with radiation exhibit normal serum biochemical parameters compared to saline treated control. Serum was collected from tumor bearing mice treated with saline, AMD3100 SPNPs, IR or AMD3100 SPNPs + IR at 23 DPI. For each treatment group levels of Creatinine, blood urea nitrogen (BUN), Alanine transaminase (ALT), Aspartate transaminase (AST), Alkaline phosphatase (ALKP), Glucose (GLUC), Total Protein (TPRO), Aluminium diboride (ALB2), Total Bilirubin (TBIL) and Calcium (Ca) were quantified. The levels of different serum biochemical parameters between the treatment groups were compared and were found non-significant, P> 0.05 (n=3 biological replicates).

**Figure S9.**
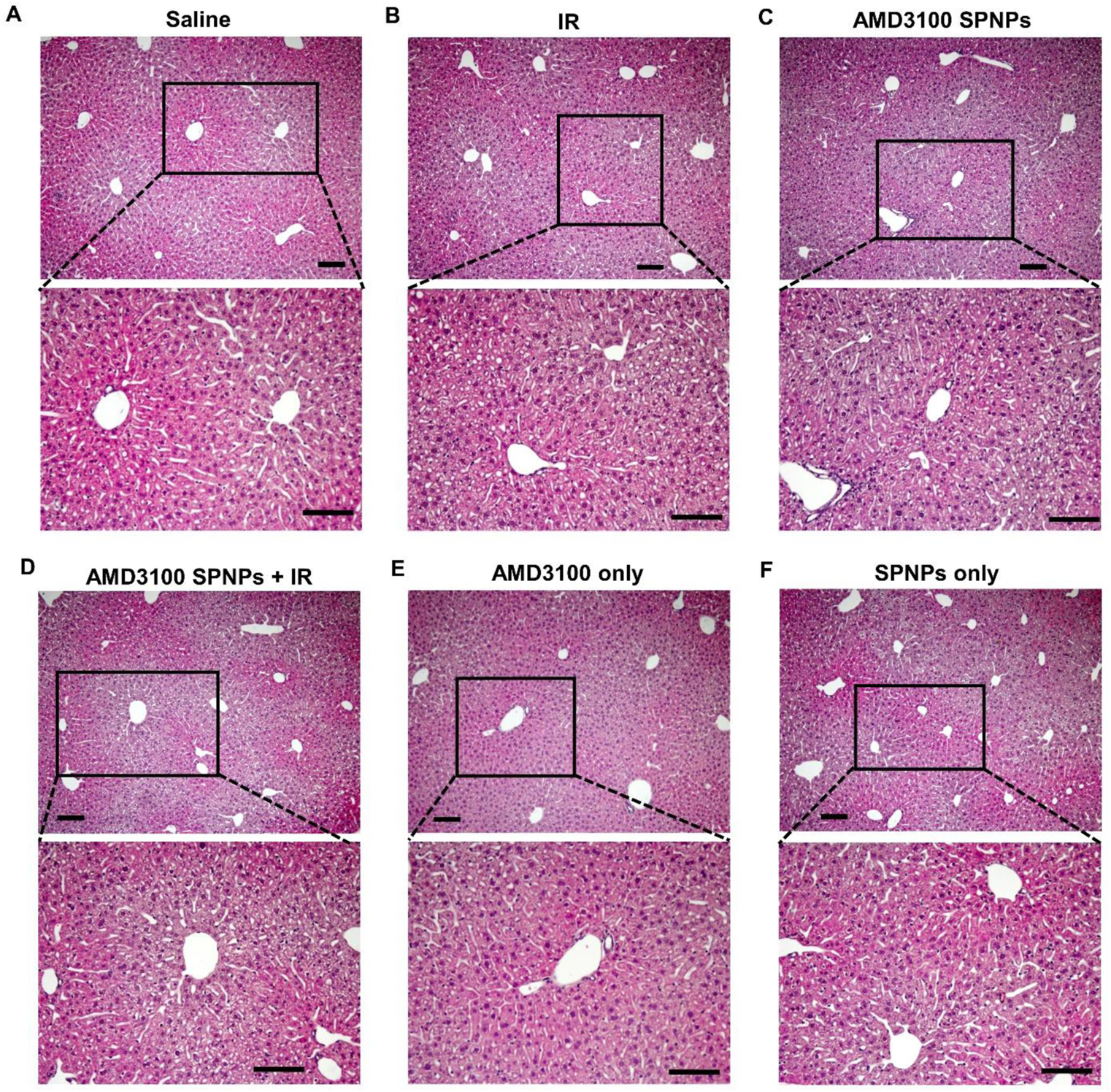
Histopathological assessment of livers from tumor bearing mice treated with AMD3100 SPNPs + IR. H&E staining of 5µm paraffin embedded liver sections from **Saline (A), IR (B)**, **AMD3100 SPNPs (C)**, **AMD3100 SPNP + IR (D)**, **AMD3100 only (E)** and **SPNPs only (F)** treatment groups. In each treatment groups, top panel represents the low magnification (10X) and bottom panel represents the high magnification (40X). High magnification panels indicate areas delineated in the low magnification panels. Histology performed on resected livers following complete treatment of OL-61 tumor bearing mice. There are no differences in the hepatocytes and the stromal central and portal areas between the control saline group and the different treatment groups. Representative images from a single experiment consisting of independent biological replicates are displayed. Black scale bars = 100µm (10X) 50µm (40X)

**Figure S10.**
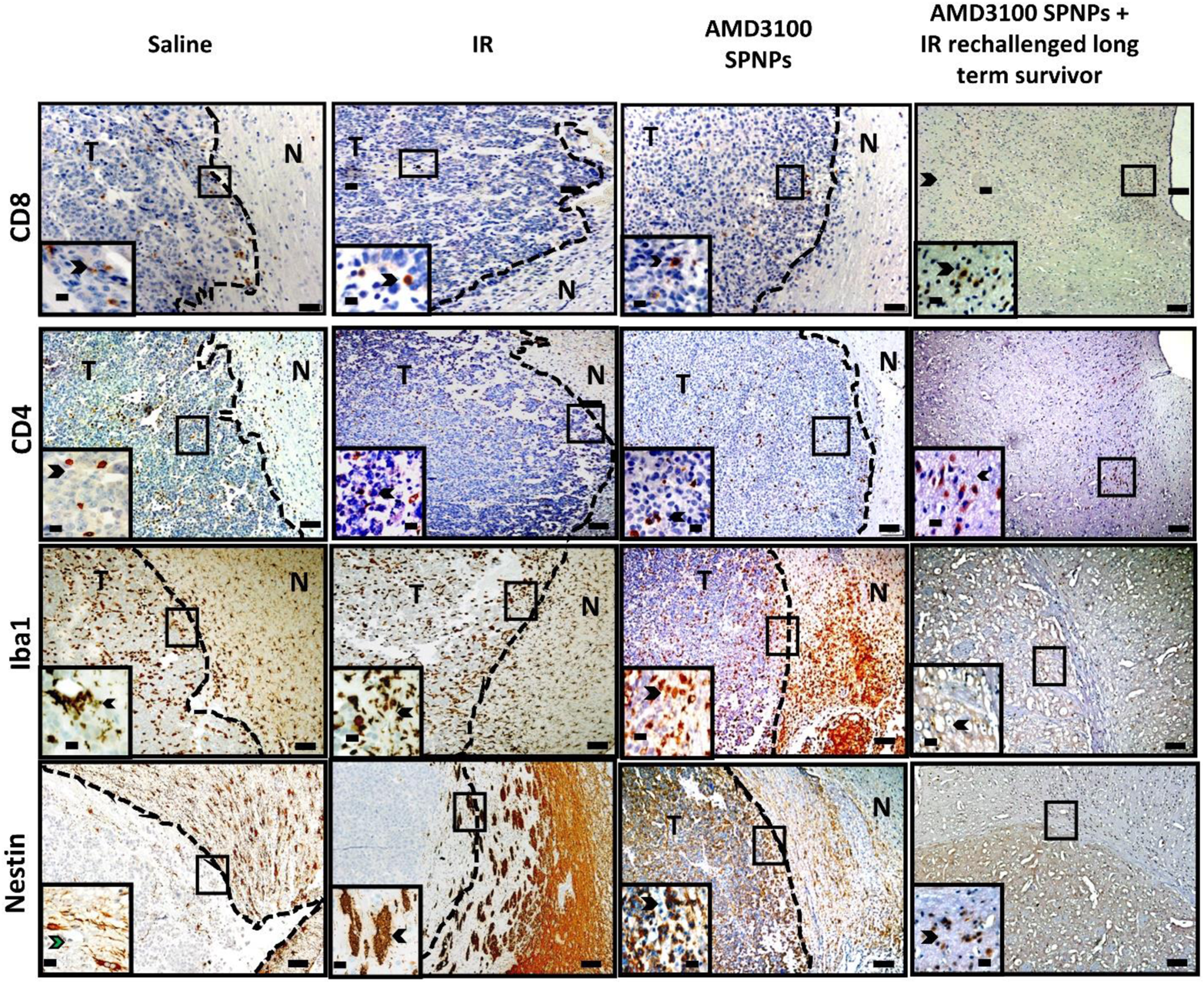
AMD3100 SPNPs treatment in combination with radiation enhances immune infiltration of glioma bearing mice. Immunohistochemistry staining of 5µm paraffin-embedded brain sections from saline (24 dpi), IR (48 dpi), AMD3100 SPNPs alone (45 dpi) and long-term survivors from AMD3100 SPNPs + IR treatment groups (60 dpi after rechallenging with OL61 cells) (scale bar = 1mm). Paraffin embedded 5µm brain sections for each treatment groups were stained for CD8, CD4, ionized calcium binding adaptor molecule 1 (Iba1) and Nestin. Low magnification (10X) panels show normal brain (N) and tumor (T) tissue (black scale bar = 100µm). Black arrows in the high magnification (40X) panels (black scale bar = 20µm) indicate positive staining for the areas delineated in the low-magnification panels. Representative images from a single experiment consisting of independent biological replicates are displayed.

**Figure S11.**
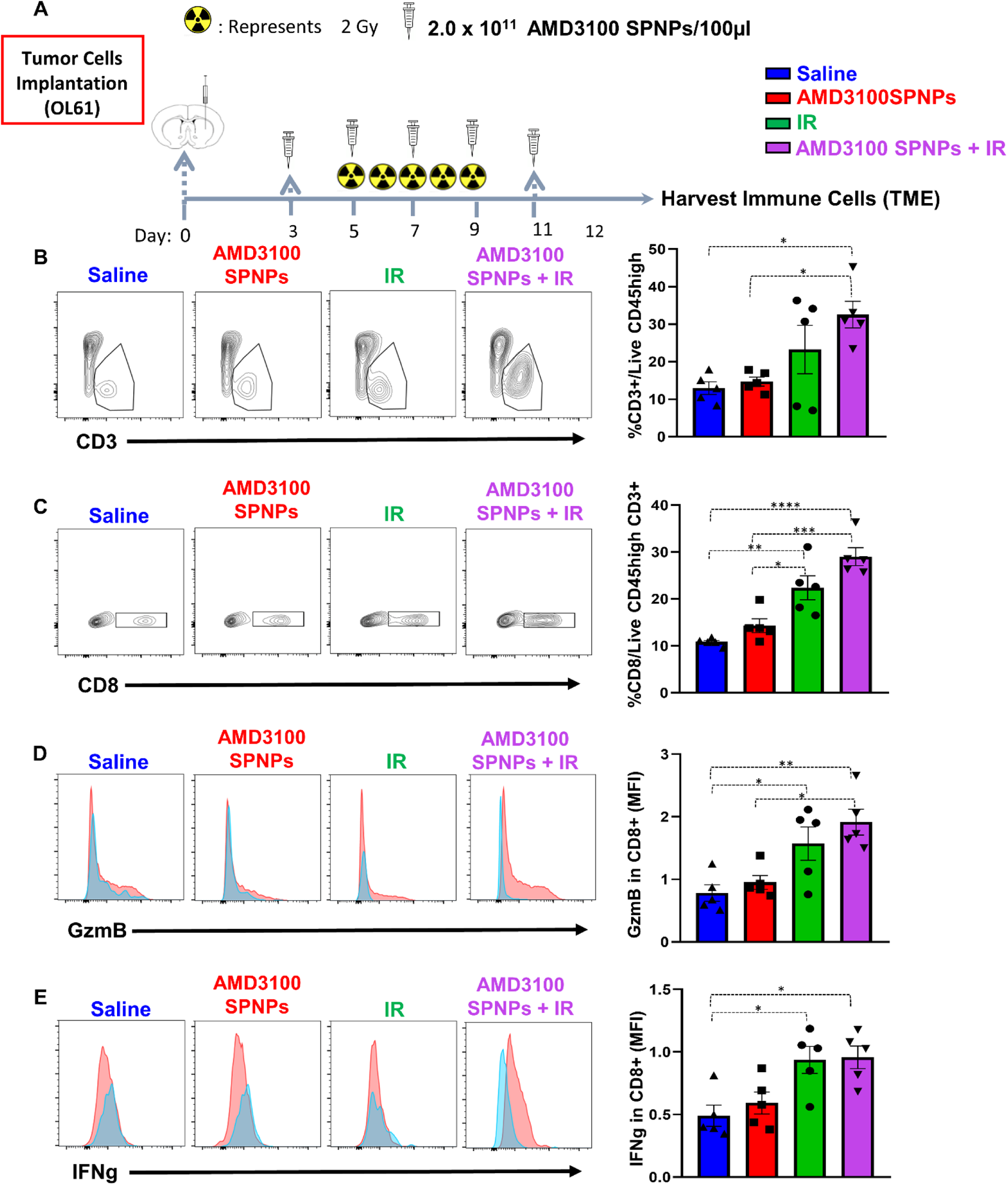
Combining AMD3100 SPNPs+ IR enhances the adaptive antitumor immune response. **(A)** Experimental design represents the timeline for the combination treatment of AMD3100 SPNPs + IR to assess the efficacy of GBM-infiltrating T cell function. **(B)** Representative flow cytometry plots and analysis represents the frequency CD3+ lymphocytes within the TME in saline, AMD 3100 SPNPs, IR or aMD3100 SPNPs+ IR group. **(C)** Representative flow cytometry plots and analysis represents the frequency CD8+ lymphocytes within the TME in saline, AMD3100 SPNPs, IR or AMD3100 SPNPs+ IR group. **(D, E)** Representative flow cytometry plots and analysis represent the expression of effector T cells molecules Granzyme B (GzmB) (C) and IFNg (D) in CD8 T cells in filtrating the TME of each group. *p< 0.05 **p< 0.01, ***p< 0.0001, ****p<0.0001; One way ANOVA. Bars represent mean ± SEM.

